# Combinatorial expression of neurexin genes regulates glomerular targeting by olfactory sensory neurons

**DOI:** 10.1101/2024.04.01.587570

**Authors:** Sung Jin Park, I-Hao Wang, Namgyu Lee, Hao-Ching Jiang, Takeshi Uemura, Kensuke Futai, Dohoon Kim, Evan Macosko, Paul Greer

## Abstract

Precise connectivity between specific neurons is essential for the formation of the complex neural circuitry necessary for executing intricate motor behaviors and higher cognitive functions. While *trans*-interactions between synaptic membrane proteins have emerged as crucial elements in orchestrating the assembly of these neural circuits, the synaptic surface proteins involved in neuronal wiring remain largely unknown. Here, using unbiased single-cell transcriptomic and mouse genetic approaches, we uncover that the neurexin family of genes enables olfactory sensory neuron (OSNs) axons to form appropriate synaptic connections with their mitral and tufted (M/T) cell synaptic partners, within the mammalian olfactory system. Neurexin isoforms are differentially expressed within distinct populations of OSNs, resulting in unique pattern of neurexin expression that is specific to each OSN type, and synergistically cooperate to regulate axonal innervation, guiding OSN axons to their designated glomeruli. This process is facilitated through the interactions of neurexins with their postsynaptic partners, including neuroligins, which have distinct expression patterns in M/T cells. Our findings suggest a novel mechanism underpinning the precise assembly of olfactory neural circuits, driven by the *trans*-interaction between neurexins and their ligands.

## Introduction

Forming precise and appropriate connections between the billions of neurons present in the mammalian brain is essential for enabling complex motor and cognitive outputs^1^. Changes in neural connectivity have been associated with neurological disorders such as autism spectrum disorder (ASD), schizophrenia, and intellectual disability, highlighting the importance of precise brain wiring for proper brain functioning^2–4^. To establish precise neuronal connections, axon guidance receptor proteins, which are localized at the growing edge of the axonal growth cone of projecting neurons, sense and interpret extracellular cues, thus directing the axon to its final destination within the brain ^5–7^. Upon successfully completing its journey, the projecting neuron forms a synaptic connection with its partner neuron(s), a process that is tightly regulated by various synaptic transmembrane proteins^8,9^. Traditionally, axon guidance and synaptic functioning were considered distinct processes, that occurred independently of one another. However, more recent research suggests that both of these processes may be coordinately regulated by *trans*-interactions of synaptic transmembrane proteins, such as teneurins and latrophilins^10–13^. Nonetheless, the specific synaptic transmembrane proteins responsible for precise neuronal assembly largely remained unidentified.

The murine olfactory system is a particularly attractive system for elucidating the mechanisms underlying neural circuit formation. Odorants are detected by the principal sensory neurons of the olfactory system known as olfactory sensory neurons (OSNs). Each OSN expresses a singular type of olfactory receptor (OR) from a repertoire of approximately 1,000 different ORs, and all OSN axons expressing the same OR type converge to an insular, anatomic structure within the mouse olfactory bulb (OB) to form a structure known as a glomerulus^14,15^. The location of a given glomerulus is highly stereotyped and is largely spatially invariant between all mice of the same species, thereby forming a glomerular map consisting of upwards of 1,000 distinct glomeruli^16^. The accurate generation of this glomerular map is thought to enable odor discrimination, pattern recognition, and the extraction of meaningful features from odorant inputs to this sensory system^17^.

Here, using single-cell RNA sequencing (scRNA-Seq) data obtained from tens of thousands of OSNs, we have conducted an unbiased analysis to identify synaptic membrane proteins, which are expressed in a differential manner across distinct types of OSNs. These candidate axon guidance receptors are highlighted by the neurexin family of genes, key presynaptic receptors involved in synaptic transmission, which are expressed in distinct patterns in different types of OSNs. Genetic ablation of neurexin isoforms disrupts the formation of the glomerular map. Single-nucleus RNA transcriptional profiling of the synaptic partners of the OSNs, the mitral and tufted (M/T) cells, reveals that neurexin *trans*-synaptic ligands are expressed in a cell type-specific expression pattern within these cells. Through genetic analysis, we find that these neurexin ligands are capable of guiding OSN axons in a neurexin-dependent manner to their appropriate glomeruli. These findings reveal a novel mechanism through which precise wiring is achieved between specific classes of neurons within the mammalian brain.

## Results

### Neurexins are differentially expressed in distinct populations of OSNs

All murine OSNs expressing the same OR project their axons to a highly stereotyped location within the OB to form a glomerulus. As each of the 1,000+ different types of OSN expresses a unique transcriptional program that is highlighted by axon guidance genes (Figure 1A), it suggests that there is an OR-dependent hard-wired genetic program that plays a crucial role in guiding the OSN axon to its appropriate location within the OB. To uncover the specific gene(s) that might be responsible for this axonal targeting, we analyzed scRNA-Seq data obtained from 20,375 OSNs to identify those genes that are most differentially expressed between different classes of OSN and thus most likely to contribute to OSN axonal pathfinding. Given the emerging role of *trans*-interactions between specific synaptic membrane proteins as crucial modulators in precise neuronal circuit assembly during development^10,11^, we first used SynGo, an evidence-based database of synaptic proteins^18^, and determined the synaptic membrane protein-encoding genes that were both highly expressed in OSNs and were differentially expressed across different types of OSN (Figure 1B). This subset of genes included several genes that had previously been reported to regulate OSN axonal pathfinding, such as neuropilins (Nrp1 and Nrp2) and protocadherins (Pcdhs)^19,20^, indicating that this approach was effective in identifying genes important for axonal guidance in the olfactory system. In parallel, we performed an analogous analysis of genes whose protein products are localized to growth cones, which revealed a smaller number of growth cone-localized genes that were highly expressed in OSNs and expressed in a highly variable manner in OSNs (Figure 1C). Only three genes appeared in both of these analyses, the protease, presenilin2 (Psen2), glycoprotein M6a (Gpm6a), and the central presynaptic organizer, neurexin1 (Nrxn1). Among these three genes, neurexin1 was chosen for further investigation for its role in olfactory wiring due to its higher variability and expression in OSNs compared to the other two.

**Figure 1.**
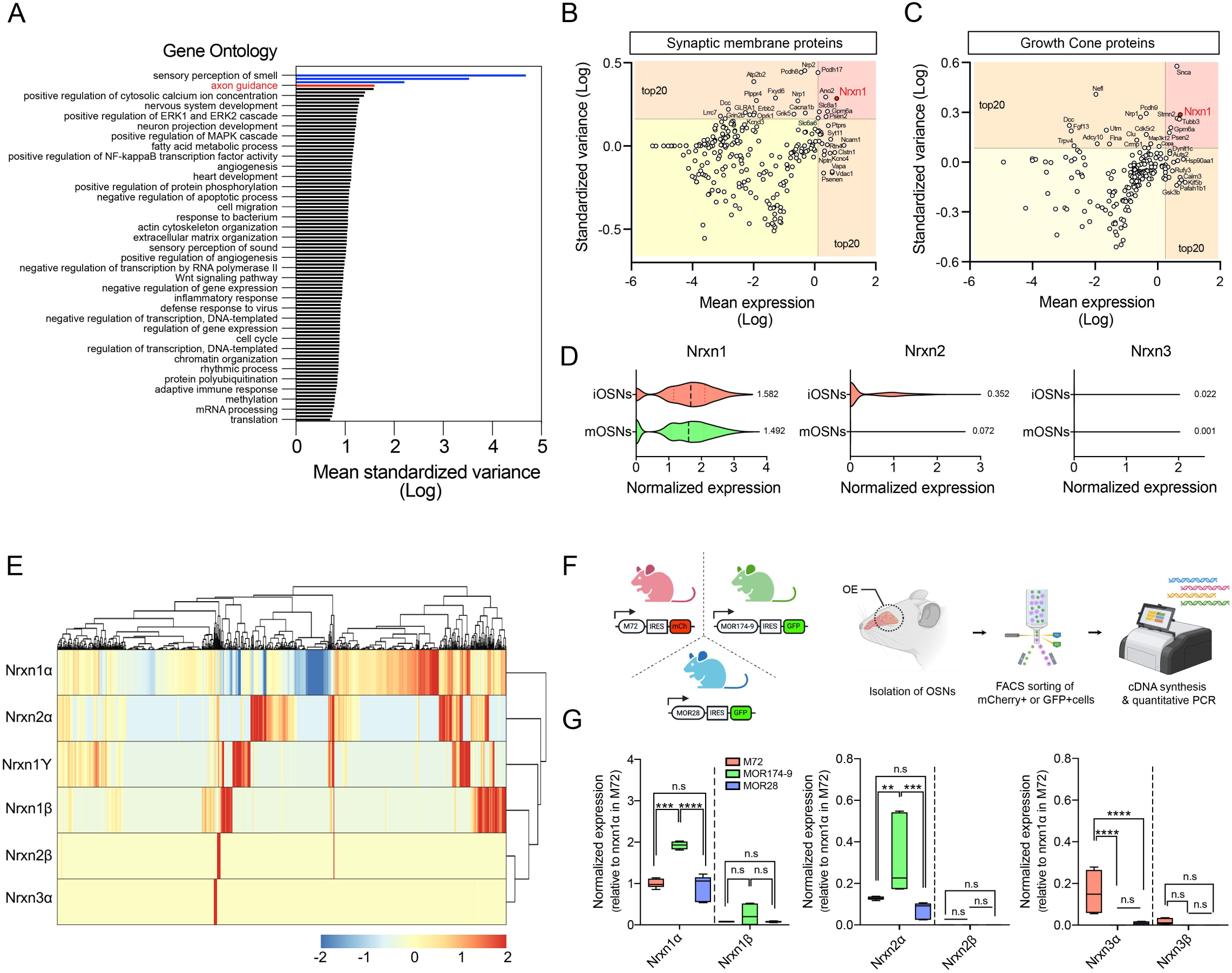
Neurexin isoforms are differentially expressed in distinct types of OSNs. (A) The differential expression of different categories of genes was determined by calculating the mean standardized variance scores of expression across OSN types. (B) Standardized variance and mean expression of 246 synaptic membrane protein encoding genes was determined. The orange boxes demarcate the 20 genes with the highest standardized variance scores or mean expression values, and the red box includes genes in the top 20 for both standardized variance and mean expression. (C) Standardized variance and mean expression of 193 genes whose protein products localize to axonal growth cones. The orange boxes demarcate the 20 genes with the highest standardized variance scores or mean expression values, and the red box includes genes in the top 20 for both standardized variance and mean expression. (D) Normalized mean expression of neurexin1, 2, and 3 (Nrxn1-3) in immature OSNs (iOSNs) and mature OSNs (mOSNs). (E) Z-score scaled heatmap showing the mean expression of neurexin alpha, beta, and gamma isoforms in OSNs types expressing 889 different ORs. (F) Schematic representation of quantitative PCR analysis of mRNAs expressed by fluorescently labeled OSNs isolated from M72-IRES-mCherry, MOR174-9-IRES-GFP, or MOR28-IRES-GFP mice. (G) Normalized expression of neurexin1, 2, and 3 alpha and beta isoforms in M72, MOR174-9, and MOR28-expressing OSNs. n = 6 for each group. **p < 0.01, ***p < 0.001, ****p < 0.0001, Tukey’s test following one-way ANOVA.

Neurexin1 is a member of a multi-gene family consisting of three different genes, neurexin1-3, which can give rise to alpha, beta, and gamma forms depending on alternative promoter usage^21–23^. Neurexin1-3 have been shown to play overlapping roles in other biological contexts, and thus to begin to investigate the potential role for neurexins in OSN axonal pathfinding, we first sought to characterize the expression patterns of all three neurexin genes in OSNs. Analysis of our single-cell transcriptomic data revealed that all three neurexin genes were highly enriched in OSNs compared to other cells whose cell bodies reside within the main olfactory epithelium of the mouse (Supplementary Figure 1). In addition, all three neurexins were expressed more highly in developing immature OSNs (iOSNs), which are actively projecting their axons to glomerular targets, compared to mature OSNs (mOSNs), whose axons have already reached their final destinations within the OB^24–26^, consistent with a potential role for neurexins in OSN wiring processes (Figure 1D).

To determine the extent to which each of the seven neurexin isoforms is differentially expressed in distinct populations of OSNs, we performed 5’ scRNA-Seq experiments, which enabled us to distinguish neurexin isoforms, which share highly similar 3’ sequences to one another. In these experiments we were able to analyze six neurexin isoforms from 889 types of OSNs that expressed different ORs from one another. An analysis of neurexin expression within these OSNs revealed a highly differential expression pattern of the neurexin isoforms across OSNs expressing different types of OR (Figure 1E).

To validate the differential expression of neurexin isoforms in OSNs, we utilized three fluorescent reporter knockin mouse lines (M72-IRES-mCherry, MOR174-9-IRES-GFP, and MOR28-IRES-GFP), which express mCherry or GFP in OSN populations expressing specific ORs (M72, MOR174-9, and MOR28, respectively) (Figure 1F). mCherry– or GFP-expressing OSNs were purified from the knockin mice using fluorescence-activated cell sorting (FACS), and mRNA was extracted for quantitative PCR analysis. These experiments revealed that neurexin1 alpha and neurexin2 alpha were most highly expressed in MOR174-9-expressing OSNs and were expressed to a lesser extent in MOR28– and M72-expressing OSNs (Figure 1G). By contrast, neurexin3 alpha exhibited significantly higher expression in M72+OSNs compared to both MOR174-9– and MOR28-expressing OSN populations. For all three of these OSN populations the beta isoforms of neurexins were expressed at much lower levels than the alpha isoforms (Figure 1G). These results are consistent with what we observed in our scRNA-Seq analyses, and together they indicate that different OSN populations exhibit a distinct expression pattern of neurexins and might participate in OSN axonal targeting.

### Neurexins are required for proper axonal targeting of OSNs

To begin to investigate the role of neurexins in the olfactory wiring, we first bred individual knockout mice for the neurexin1, 2, or 3 alpha isoforms^27^ with M72-IRES-mCherry mice to enable us to visualize glomeruli formed by OSNs expressing these ORs in the presence or absence of neurexin genes (Figure 2A). M72 was chosen for these initial experiments as our single-cell analysis had revealed that neurexin1, 2, and 3 were all highly expressed in M72-expressing OSNs (Figure 1G). Immunohistochemical analysis revealed that the ablation of any of the neurexin1, 2, or 3 alpha isoforms led to abnormal axonal coalescence of M72-expressing OSNs within the OB, resulting in a significant number of M72-expressing OSN axonal mistargeting events and the formation of additional glomeruli (Figure 2B). To extend this analysis beyond OSNs expressing M72, we next examined projections from MOR174-9-expressing OSNs and MOR28-expressing OSNs lacking neurexins using MOR174-IRES-GFP and MOR28-IRES-GFP mice, respectively (Figure 2A). These lines were selected as our transcriptional profiling experiments had revealed that only the alpha isoforms of neurexin1 and neurexin2, but not neurexin3, were expressed within these OSNs (Figure 1G). Consistent with this expression pattern, we observed that knockout of neurexin1 alpha or neurexin2 alpha, but not neurexin3 alpha, resulted in the abnormal formation of MOR174-9+ and MOR28+ glomeruli within the OB (Figure 2C and 2D). The lack of significant changes in the positioning of MOR174-9+ and MOR28+ glomeruli in neurexin3 alpha knockout mice might be attributed to the very low expression of neurexin3 alpha in these OSNs. These findings suggest that the neurexin alpha isoforms are required for OSN axonal targeting to specific glomeruli and that the contribution of individual neurexin alpha isoforms in axonal targeting for a particular type of OSN correlates with its degree of expression.

**Figure 2.**
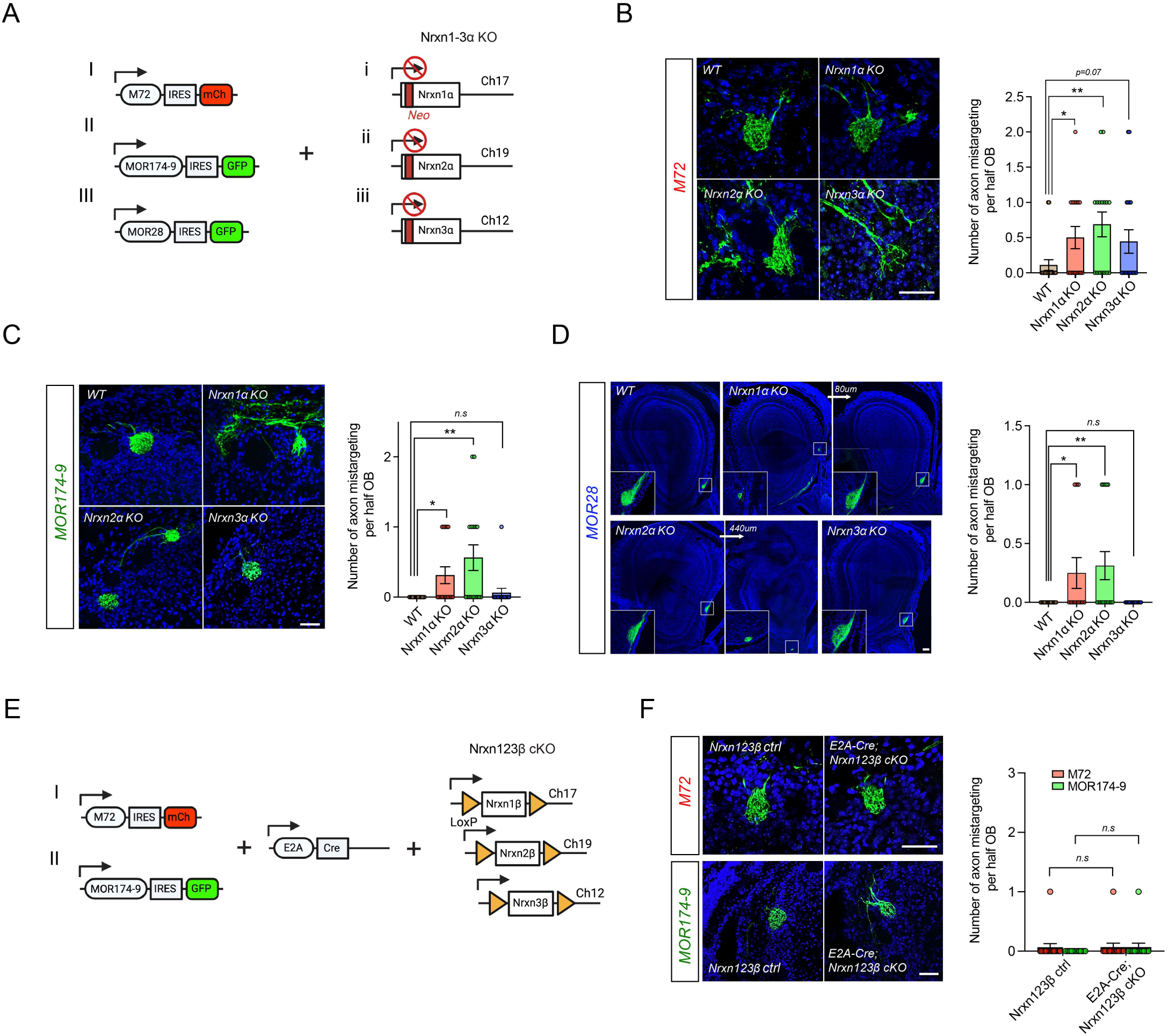
Neurexin alpha isoforms are required for precise axonal targeting of M72-, MOR174-9-, and MOR28-expressing OSNs. (A) Generation of mouse models to analyze the effect of neurexin alpha knockout (KO) on OSN axonal targeting. M72-IRES-mCherry, MOR174-9-IRES-GFP, and MOR28-IRES-GFP mice were crossed with neurexin1, 2, and 3 alpha single KO mice (Nrxn1-3α KO). (B) Representative images showing axonal targeting of M72-expressing OSNs within specific glomeruli in the OBs of wild-type (WT) and neurexin alpha KO mice (left panel), and quantification of the number of axonal mistargeting events per half OB (right panel). n = 18 for WT, 16 for Nrxn1α KO, 16 for Nrxn2α KO, 18 for Nrxn3α KO. Data are presented as mean ± SEM, *p < 0.05, **p < 0.01, Fisher’s LSD test following one-way ANOVA. Scale bar, 40 μm. (C) Representative images depicting axonal targeting of MOR174-9-expressing OSNs within specific glomeruli in the OBs of WT and neurexin alpha KO mice (left panel), and quantification of the number of axonal mistargeting events per half OB (right panel). n = 18 for WT, 16 for Nrxn1α KO, 16 for Nrxn2α KO, 16 for Nrxn3α KO. Data are presented as mean ± SEM, *p < 0.05, **p < 0.01, Fisher’s LSD test following one-way ANOVA. Scale bar, 40 μm. (D) Representative images depicting axonal targeting of MOR28-expressing OSNs within specific glomeruli in the OBs of WT and alpha neurexin KO mice (left panel), and quantification of the number of axonal mistargeting events per half OB (right panel). n = 16 for WT, 12 for Nrxn1α KO, 16 for Nrxn2α KO, 16 for Nrxn3α KO. Data are presented as mean ± SEM, *p < 0.05, **p < 0.01, Fisher’s LSD test following one-way ANOVA. Scale bar, 100 μm. (E) Generation of mouse models to analyze the effect of deleting neurexin beta genes on OSN axonal targeting. M72-IRES-mCherry and MOR174-9-IRES-GFP mice were crossed with E2A-Cre and triple neurexin beta floxed mice (Nrxn123β cKO). (F) Representative images showing axonal targeting of M72– or MOR174-9-expressing OSNs within specific glomeruli in the OBs of triple neurexin beta control (ctrl) and E2A-Cre-dependent triple neurexins beta conditional KO (cKO) mice (left panel). Quantification of these analyses of axonal mistargeting is shown in the right panel. n = 16 for Nrxn123β ctrl in M72, 15 for E2A-Cre;Nrxn123β cKO in M72, 14 for Nrxn123β ctrl in MOR174-9, 15 for E2A-Cre;Nrxn123β cKO in MOR174-9. Data are presented as mean ± SEM, Fisher’s LSD test following one-way ANOVA. Scale bar, 40 μm.

If this hypothesis was correct, we would predict that deletion of the beta isoforms of neurexin would have minimal effect on the axonal targeting of these three OSN populations as the beta isoforms of all three neurexins were barely detected within these cells (Figure 1G). To test this, we utilized mice in which all three beta isoforms of neurexin were flanked by loxP sites^28^, facilitating the constitutive knockout of these genes when bred to E2A-Cre mice, which express Cre recombinase throughout the body^29^ (Figure 2E). In contrast to the alpha neurexin individual knockout mice, the position and number of M72+ or MOR174-9+ glomeruli remained unchanged in mice in which all three neurexin beta isoforms were deleted (Figure 2F). Importantly, neither the overall structure and size of the OB nor the total number of glomeruli were affected in neurexin knockout mice (Supplementary Figure 2A-2C), indicating that deletion of neurexin genes specifically disrupts OSN axonal projections rather than affecting the overall architecture of the OB.

### OSN-specific neurexin isoforms synergistically control OSN axonal targeting

The experiments described above all involved the constitutive knockout of neurexin isoforms throughout all cells of the olfactory system. Our scRNA-Seq analysis had revealed that neurexins are expressed within OSNs (Figure 1D), but additional single-nucleus RNA sequencing (snRNA-Seq) experiments also indicated that neurexin genes are also expressed within M/T cells, the synaptic partners of the OSNs (Figure 3A), thus raising the possibility that neurexins could be acting either within OSNs or M/T cells to regulate glomerular formation. To distinguish between these possibilities, we utilized conditional neurexin1 knockout mice^30^, in which M72-expressing OSNs were labeled with mCherry in combination with three different Cre mice (Figure 3B): E2A-Cre, which expresses Cre recombinase in all cells of the body, Goofy-Cre, which leads to Cre expression specifically in postmitotic OSNs^31^, and Pcdh21-Cre, which expresses Cre in postmitotic M/T cells^32^, to enable us to determine the effect of deleting neurexin1 from different olfactory cell populations on M72+ glomerular formation^31,32^. Conditional deletion of neurexin1 from either all cells or from OSNs phenocopied the disruption in M72+ glomeruli observed with constitutive deletion of neurexin1 (Figure 3C). By contrast, M/T cell-specific neurexin1 deletion resulted in normal axonal targeting of M72+OSNs (Figure 3C). These findings indicate that the neurexin acts within OSNs to regulate glomerular targeting.

**Figure 3.**
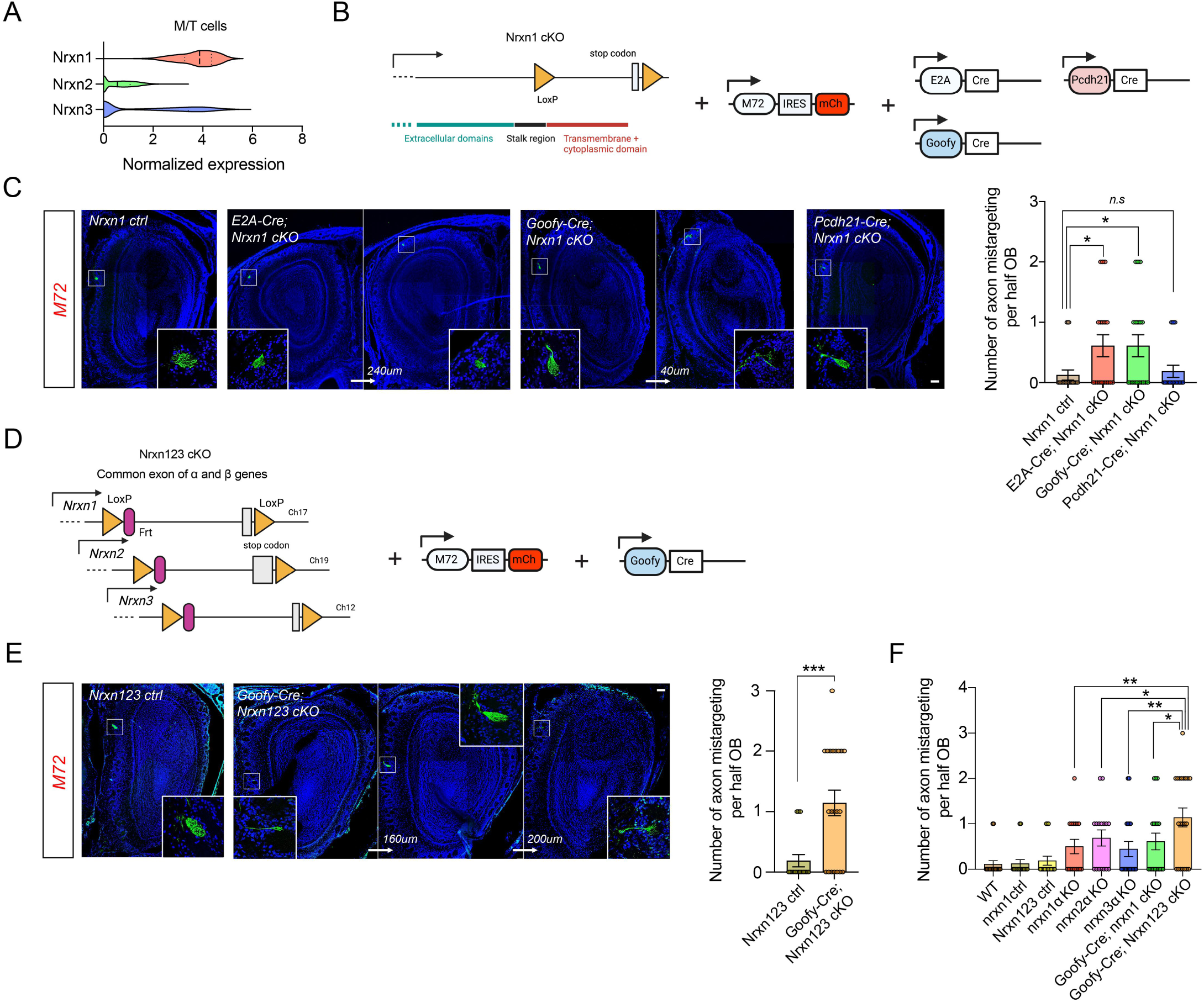
OSN-specific neurexin isoforms cooperatively govern OSN axonal targeting. (A) Neurexin1, 2, and 3 (Nrxn1-3) normalized mean expression in M/T cells. (B) Generation of mouse models for analysis of the effect of cell type-specific neurexin1 conditional knockout (cKO) on OSN axonal targeting. M72-IRES-mCherry mice were bred with E2A-Cre, Goofy-Cre, or Pcdh21-Cre mice and further crossed with neurexin1 floxed mice (Nrxn1 cKO). (C) Representative images showing axonal targeting of M72-expressing OSNs within specific glomeruli in the OBs of wild-type (Nrxn1 ctrl) and E2A-Cre-, Goofy-Cre-, and Pcdh21-Cre-dependent neurexin1 cKO mice (left panel). Quantification of axonal mistargeting is shown in the right panel. n = 16 for Nrxn1 ctrl, 18 for E2A-Cre;Nrxn1 cKO, 18 for Goofy-Cre;Nrxn1 cKO, 16 for Pcdh21-Cre;Nrxn1 cKO. All data were collected on at least three separate days. Data are presented as mean ± SEM, *p < 0.05, Fisher’s LSD test following one-way ANOVA. Scale bar, 100 μm. (D) Schematic of the generation of mouse models for an analysis of OSN-specific triple neurexin cKO effects on OSN axonal targeting. M72-IRES-mCherry mice were crossed with Goofy-Cre and triple neurexin floxed mice (Nrxn123 cKO). (E) Representative images showing axonal targeting of M72-expressing OSNs within specific glomeruli in the OBs of wild-type (Nrxn123 ctrl) and Goofy-Cre-dependent triple neurexin cKO mice (left panel), with statistical analysis of axonal mistargeting (right panel). n = 22 for Nrxn123 ctrl, 21 for Goofy-Cre;Nrxn123 cKO. All data were collected on at least three independent days. Data are presented as mean ± SEM, ***p < 0.001, unpaired t test. Scale bar, 100 μm. (F) Quantification of the number of axonal mistargeting per events for neurexin single KO (Nrxn1-3α KO), OSN-specific neurexin1 cKO (Goofy-Cre;Nrxn1 cKO), and OSN-specific triple neurexin cKO (Goofy-Cre;Nrxn123 cKO) mice from figures 2B, 3C, and 3E, respectively. Data are presented as mean ± SEM, *p < 0.05, **p < 0.01, Fisher’s LSD test following one-way ANOVA.

All of the experiments performed to this point were done using mouse lines in which individual neurexin alpha isoforms were deleted. However, our transcriptomic profiling revealed that many OSN types co-expressed multiple neurexin genes. To determine whether neurexins interacted synergistically to regulate OSN axonal projections, we used a mouse line in which loxP sequences flank the common last exon of each alpha neurexin and beta neurexin gene, allowing for conditional knockout of all neurexin isoforms^33^. These mice were crossed with M72-IRES-mCherry mice and Goofy-Cre or Pcdh21-Cre mice (Figure 3D). OSN-specific knockout of all three neurexin genes led to a significant increase in the number of misprojected M72+OSN axons compared to control and other single knockout mice in which individual neurexin alpha isoforms were deleted (Figure 3E and 3F), suggesting that neurexins synergistically cooperate to regulate OSN axonal targeting to specific glomeruli.

### Neurexin ligands have distinct expression patterns in M/T cell subtypes

The above experiments reveal that neurexins function within OSNs to regulate the guidance of OSN axons to appropriate glomeruli within the OB where they can form synaptic connections with M/T cells. Neurexins have been shown to form *trans*-synaptic complexes with a number of different postsynaptic ligands, including neuroligins, neurexophilins, and leucin-rich repeat transmembrane protein2 (LRRTM2), and these interactions are essential for neurexin’s ability to regulate synaptic transmission^34^. Based on these observations, we hypothesized that neurexins expressed at OSN axonal termini interact with ligands expressed on the dendrites of M/T cells to regulate appropriate wiring of OSNs with M/T cells within the OB. To begin to test this hypothesis, we first examined the expression patterns of neurexin ligands in M/T cells using snRNA-Seq to determine which neurexin ligands are expressed by M/T cells. Transcriptomic profiling of 10,599 unique M/T nuclei revealed significant expression of multiple genes encoding neurexin ligands. Of these neurexin ligands, neuroligin1 exhibited the highest average expression level within M/T cells (Figure 4A).

**Figure 4.**
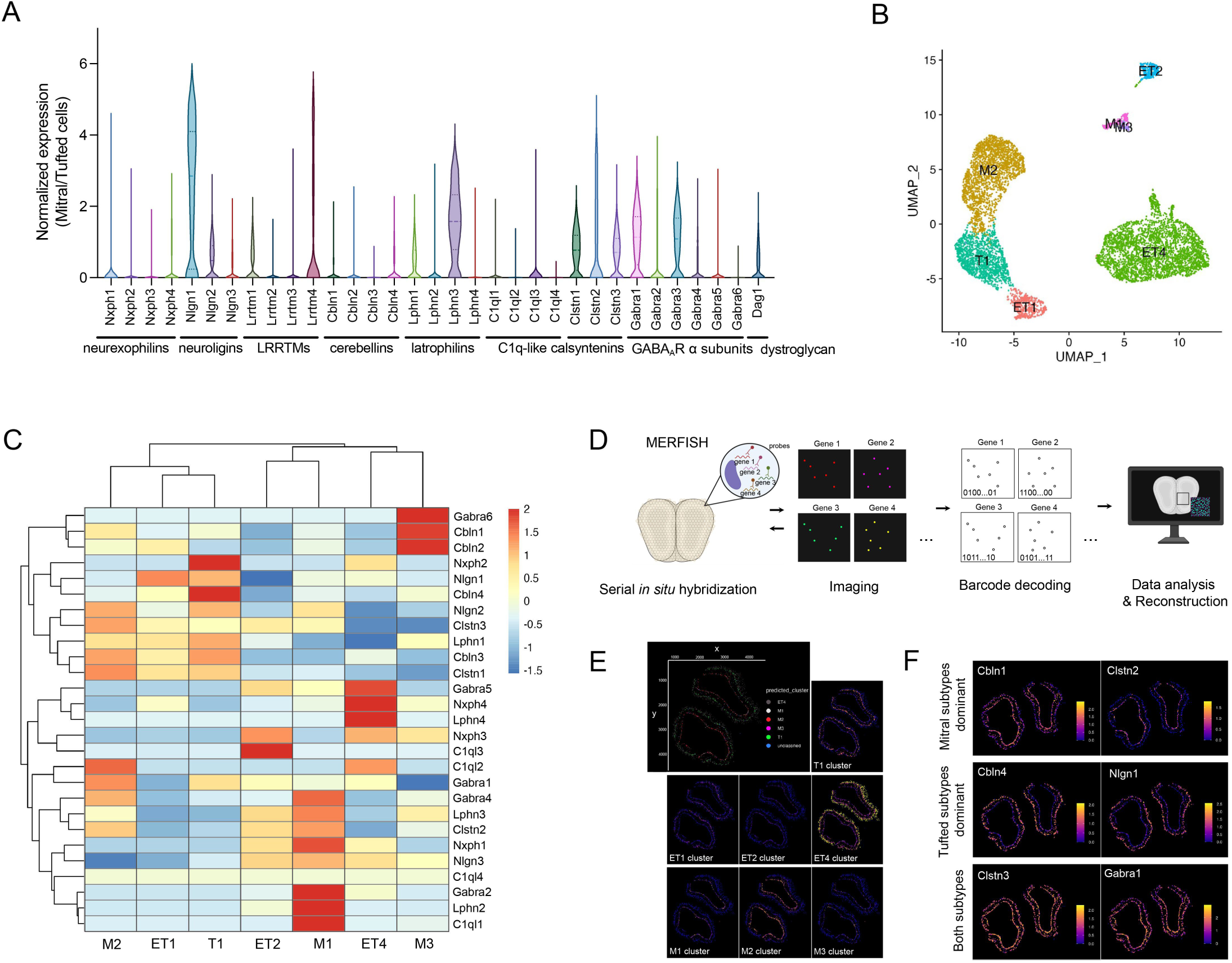
M/T cells have distinct expression patterns of neurexin ligands. (A) Normalized mean expression of neurexin postsynaptic partners in M/T cells. (B) UMAP plot revealing the different M/T cell subtypes found within the OB. M1, M2, and M3 are mitral cell subtypes, and ET1, ET2, ET4, and T1 are tufted cell subtypes. (C) Z-score scaled heatmap describing the expression patterns of neurexin postsynaptic ligands in M/T cell subtypes. (D) Schematic representation of multiplexed error-robust fluorescence in situ hybridization (MERFISH) for spatially visualizing the expression of neurexin postsynaptic partner mRNAs in M/T cells within the OB. (E) Identification of M/T cell subtypes within the OB using M/T cell subtype-specific marker genes. (F) M/T cell subtype-specific expression of neurexin postsynaptic ligands.

To examine the expression patterns of neurexin ligands within M/T cells in greater detail, we subclustered M/T nuclei using marker genes previously employed in M/T single cell profiling^35^. This procedure resulted in the identification of three distinct subtypes of mitral cells (M1, M2, M3) and four subtypes of tufted cells (ET1, ET2, ET4, and T1) which have all been described previously^35^ (Figure 4B). Intriguingly, the expression of neurexin ligands within M/T cell subtypes was not homogeneous, indicating that similarly to the neurexins themselves, neurexin ligand genes were also differentially expressed across distinct populations of M/T cells (Figure 4C).

To obtain higher resolution information about the expression patterns of neurexin ligands in M/T cells, we employed multiplexed error-robust fluorescence in situ hybridization (MERFISH), a technique that enables quantitative and spatial analysis of RNA expression at single cell resolution^36^ (Figure 4D). These experiments aligned with the snRNA-Seq analysis and revealed that while some neurexin ligands, including calsyntenin3 and type A GABA receptor subunit1 (Gabra1), were broadly expressed across multiple M/T cell types, others, such as cerebellin1 and calsyntenin2, were predominantly detected in mitral cell types, and cerebellin4 and neuroligin1 were predominantly expressed in tufted cell types (Figure 4E, 4F, and Supplementary Figure 3). Together, these experiments reveal that, much like the neurexin genes themselves, neurexin ligands are also expressed differentially across M/T cellular populations, suggesting that they might cooperate with neurexin genes to regulate the appropriate connections between OSNs and M/T cells.

### A gradient of neurexin ligands induces axon attraction *in vitro* in a neurexin-dependent manner

To begin to investigate the potential role of neurexin ligands in guiding neurexins-expressing neuronal axons, we took advantage of a microfluidic based *in vitro* axonal isolation chamber device, which consists of segregated somal and axonal compartments, which are interconnected by parallel microchannels^37^ (Figure 5A). With this device, we established gradients of soluble purified neurexin ligands in the axonal compartment and assessed the effect of these proteins on axonal attraction or repulsion^37,38^. Cortical neurons, which express high levels of neurexins, particularly neurexin1 alpha (Supplementary Figure 4A), and which are commonly employed for *in vitro* axon guidance studies^39,40^, were utilized for these experiments owing to the difficulty of culturing OSNs^41,42^. Gradients of purified, soluble neurexin ligands, neurexophilin1, LRRTM2, latrophilin1, calsyntenin3, and neuroligin1^43–48^ were all tested for their effect on axonal pathfinding using this microfluidic based assay (Figure 5B). Each of these purified recombinant proteins was able to efficiently bind to neurexin1 alpha-expressing HEK293 cells (Supplementary Figure 4B and 4C). Quantitative analysis of axons in the axonal compartment revealed a significant increase in axonal elongation in response to gradients of latrophilin1, calsyntenin3, and neuroligin1 recombinant proteins; by contrast, a gradient of LRRTM2 reduced axonal growth (Figure 5B and 5C). To assess whether the effect of neurexin ligands was merely a result of general trophic effect on axonal outgrowth, recombinant neurexin ligands were also directly added to cortical neurons cultured in standard round well plates such that a homogeneous concentration of these ligands was established. When delivered to neurexin-expressing neurons in this manner such that no gradient of ligand is present, only recombinant LRRTM2 proteins significantly altered neurite length, indicating that the alteration in axonal elongation observed in the presence of a gradient of LRRTM2 recombinant protein might be attributed to the neurotrophic effect of this ligand in inhibiting neurite outgrowth rather than a gradient-specific axonal guidance effect (Supplementary Figure 4D and 4E). Together, these findings demonstrate that gradients of latrophilin1, calsyntenin3, and neuroligin1 serve as cues for axon attraction without affecting overall axonal growth.

**Figure 5.**
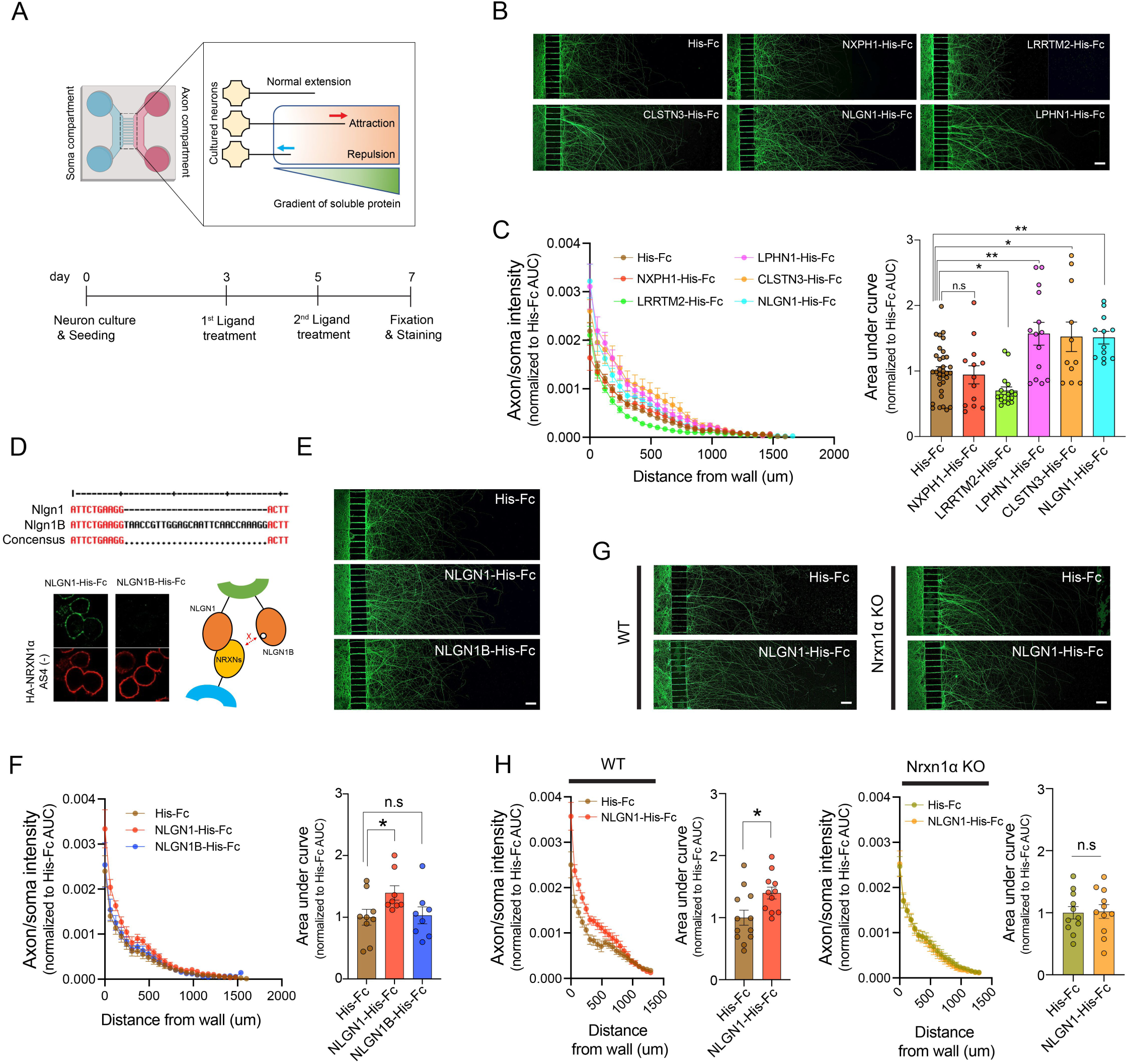
Gradients of neurexin ligands induce neurexin-dependent axonal attraction *in vitro*. (A) Schematic of the *in vitro* microfluidic based axonal isolation chamber assay. Dissociated embryonic cortical neurons were cultured in the “Soma” compartment for three days. Axons in the axonal compartment were exposed to gradients of His-tagged recombinant proteins to analyze their effect on axonal attraction or repulsion. (B) Representative images of axons exposed to various neurexin ligand recombinant proteins. NLGN1; neuroligin1, LPHN2; latrophilin2, LRRTM2; leucin-rich repeat transmembrane protein2, NXPH1; neurexophilin1, CLSTN3; calsyntenin3. Scale bar, 100 μm. (C) Quantitative analysis of axonal fluorescence intensities at different distances from the wall following exposure to gradients of different neurexin ligands (left panel). Statistical comparison of area under the curve (AUC) of the graphs from the left panel (right panel). Fluorescence intensities in the axonal compartment were normalized to those in the somal compartments and further normalized to AUC values of fluorescence graphs from the His-Fc control group, (which was set at the value of 1). n = 33 for His-Fc, 13 for NXPH1-His-Fc, 17 for LRRTM2-His-Fc, 14 for LPHN2-His-Fc, 12 for NLGN1-His-Fc, 11 for CLSTN3-His-Fc. All data were collected from at least three independent days. Data are presented as mean ± SEM, *p < 0.05, **p < 0.01, Fisher’s LSD test following one-way ANOVA. (D) Nucleic acid sequence differences between neuroligin1 and neuroligin1B (top panel), leading to reduced binding affinity between neurexin and neuroligin1B (bottom panel). (E) Representative images of axons exposed to neuroligin1 and neuroligin1B recombinant proteins. Scale bar, 100 μm. (F) Quantitative analysis of axonal fluorescence intensities upon exposure to recombinant neuroligin1 proteins (left panel). Statistical comparison of area under the curve (AUC) of the graphs from the left panel (right panel). Fluorescence intensities in the axonal compartment were normalized to those in the somal compartments and further normalized to AUC values of fluorescence graphs from the His-Fc control group. n = 9 for His-Fc, 8 for NLGN1-His-Fc, 8 for NLGN1B-His-Fc. All data were collected from three independent days. Data are presented as mean ± SEM, *p < 0.05, Fisher’s LSD test following one-way ANOVA. (G) Representative images of wild-type (WT) and neurexin1 alpha knockout (Nrxn1α KO) axons exposed to control or neuroligin1 recombinant proteins. Scale bar, 100 μm. (H) Quantitative analysis of axonal fluorescence intensities in response to exposure to gradients of neuroligin1 recombinant proteins in WT (left panels) and Nrxn1α KO neurons (right panels). Fluorescence intensities in the axonal compartment were normalized to those in the somal compartments and further normalized to AUC values of fluorescence graphs from the His-Fc control group. n = 12 for His-Fc in WT, 11 for NLGN1-His-Fc in WT, 11 for His-Fc in Nrxn1α KO, 11 for NLGN1-His-Fc in Nrxn1α KO. All data were collected on at least three independent days. Data are presented as mean ± SEM, *p < 0.05, unpaired t test.

Of these three neurexin ligands, neuroligin1, was selected for further study as it is the best characterized neurexin postsynaptic partner and is expressed at the highest levels in M/T cells. To begin to investigate whether the effect of neuroligin1 on axonal outgrowth occurred in a neurexin-dependent manner, we first employed a splicing variant of neuroligin1, which contains a splice site B (neuroligin1B) resulting in this protein having a low binding affinity for neurexins^47^ (Figure 5D). Consistent with neuroligin1 modulating axonal outgrowth in a neurexin-dependent manner, neuroligin1B had no effect on axonal growth in neurexins-expressing neurons (Figure 5E and 5F). To extend these observations, neuronal cultures from wild-type or neurexin1 alpha knockout mice were exposed to a gradient of neuroligin1 recombinant protein. Neurexin1 alpha knockout completely abolished the attractive effect of neuroligin1, indicating that neuroligin1 regulates axonal outgrowth in a neurexin-dependent manner (Figure 5G and 5H).

### M/T cell-specific neuroligin1 knockout leads to OSN axonal mistargeting

While intriguing, the effect of neuroligin1 on neurexin-mediated outgrowth was observed *in vitro* and in cortical neurons rather than OSNs. To determine more definitely whether neuroligin was important for neurexin-mediated OSN axonal pathfinding *in vivo*, we employed neuroligin1 floxed mice, which carry two loxP sequences flanking exon7 of the neuroligin1 gene, allowing for conditional knockout of neuroligin1^49^ (Figure 6A). These neuroligin1 floxed mice were crossed with M72-IRES-mCherry mice and three different Cre mice (E2A-Cre, Goofy-Cre, and Pcdh21-Cre) (Figure 6A), and we subsequently monitored the axonal targeting of M72+OSNs. Deletion of neuroligin1 in all cells through the use of E2A-Cre or specifically in M/T cells using Pcdh21-Cre mice led to incorrect axonal targeting of M72+OSNs that mimicked what we had observed in neurexin knockout mice (Figure 6B). By contrast, no deficit in M72 glomerular targeting was observed when neuroligin1 was knocked out specifically in OSNs (Figure 6B), suggesting that neuroligin1 expressed within M/T cells regulates OSN axonal targeting through its *trans*-synaptic interactions with neurexins.

**Figure 6.**
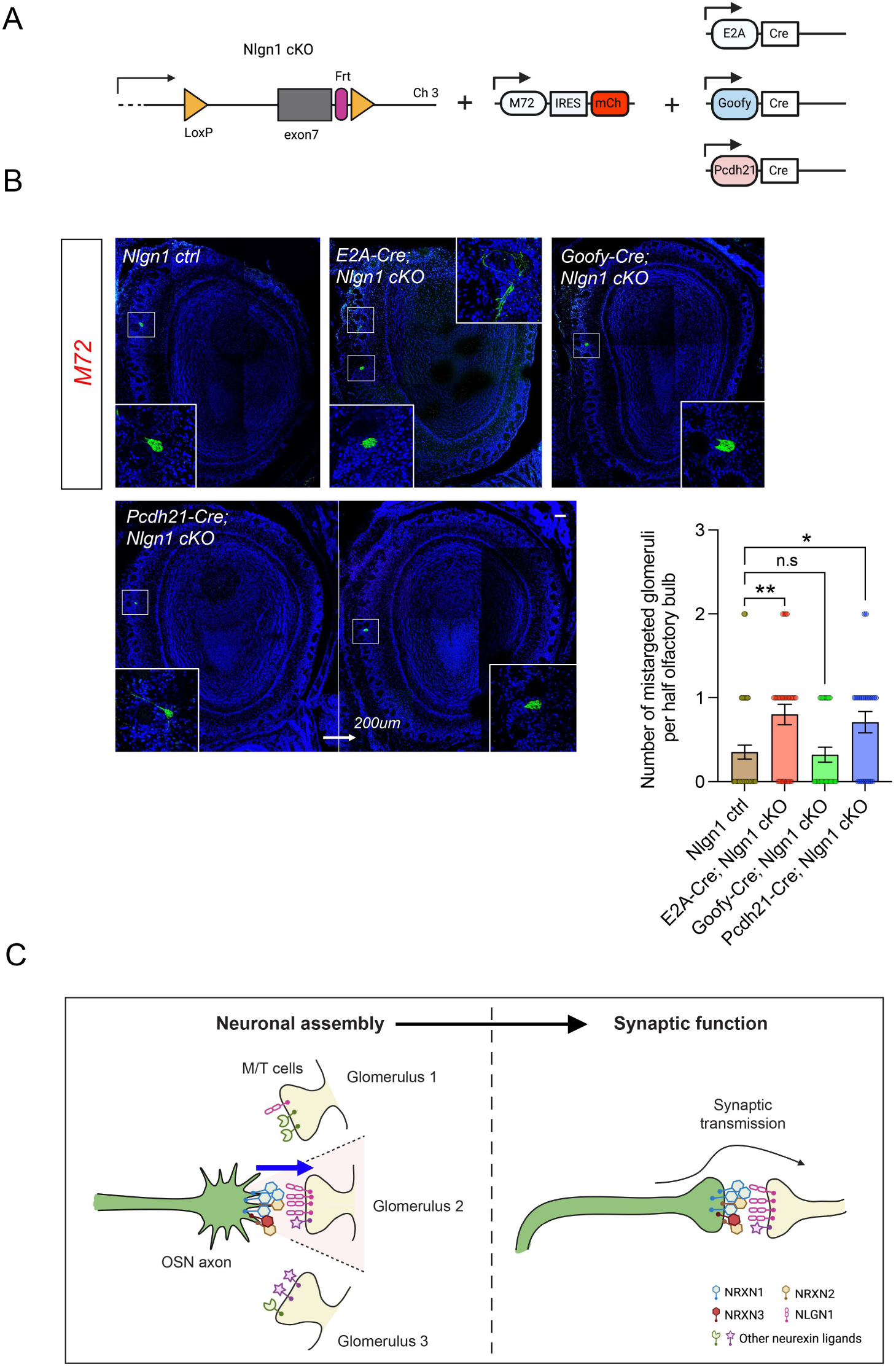
M/T cell-specific neuroligin1 knockout causes OSN axonal mistargeting. (A) Schematic of the generation of mouse models for analyzing cell type-specific neuroligin1 conditional knockout (cKO) effects on OSN axonal targeting. M72-IRES-mCherry mice were bred to E2A-Cre, Goofy-Cre, or Pcdh21-Cre mice and further crossed with neuroligin1 floxed mice (Nlgn1 cKO). (B) Representative images illustrating axonal targeting of M72-expressing OSNs within specific glomeruli in the OBs of neuroligin1 control (ctrl) and E2A-Cre-, Goofy-Cre-, and Pcdh21-Cre-dependent neuroligiin1 cKO mice (left panel), with statistical analysis of the number of axonal mistargeting events per half OB (right panel). n = 29 for Nlgn1 ctrl, 30 for E2A-Cre;Nlgn1 cKO, 28 for Goofy-Cre;Nlgn1 cKO, 24 for Pcdh21-Cre;Nlgn1 cKO. All data were collected on at least three independent days. Data are presented as mean ± SEM, *p < 0.05, **p < 0.01, Fisher’s LSD test following one-way ANOVA. Scale bar, 100 μm. (C) A schematic diagram illustrating the dual role of *trans*-interactions between neurexins and their partners in axon guidance during early development and synaptic transmission in mature synapses within the olfactory system.

## Discussion

Achieving precise synaptic connections between specific neurons is essential for organisms to be able to properly execute cognitive and motor functions. This problem is especially daunting within the mammalian brain, which consists of billions of neurons that form trillions of synapses with one another. The challenges faced by the developing mammalian nervous system are highlighted within the mouse olfactory system, which possesses upwards of one thousand distinct OSN subtypes, each of which must guide its axon to a stereotyped location within the mouse OB to form synaptic connections with their partner M/T cells. The precise pairing of OSNs with M/T cells forms the basic computational unit of the olfactory system and disruption of this wiring interferes with the ability of the olfactory system to discriminate odorants and to respond appropriately to them. Considerable effort has gone into understanding the mechanisms by which OSN axons are guided to their appropriate glomeruli, which has provided significant insight into this process, but our understanding of how OSNs form appropriate connections with their partner M/T cells remains poorly understood. Here, we identify a crucial role for the neurexin family of genes in the ability of OSNs to correctly form glomerular structures. Genetic deletion of neurexin genes from OSNs or deletion of their *trans*-synaptic ligands, the neuroligins, from M/T cells disrupts the ability of OSNs to correctly navigate to their appropriate glomerular location and form synaptic connections with their M/T cell partners.

The identification of a critical role for neurexin genes in axonal pathfinding is especially interesting considering that the vast majority of prior work on neurexins has focused on their role in synaptic transmission in the mature brain^34^ (Figure 6C). Our findings thus add to a small, but growing body of literature, which has found that proteins such as teneurin and latrophilin play dual roles in the establishment of synaptic connections as well as the subsequent functioning of the resultant synapses. The coordinate regulation of synaptic pairing and function makes sense from a conceptual perspective, ensuring a well-orchestrated process. Nonetheless, how the same gene mediates both of these disparate processes remains unclear. It is intriguing to speculate that perhaps either the identity of the *trans*-synaptic ligand changes during development or new cis-interacting protein partners emerge during synaptic development resulting in an alteration in downstream signaling events, but future experiments will be required to elucidate how neurexins are able to accomplish these dual functions. It is important to note that recent studies outside of the mammal using in vitro assays and model organisms such as fruit flies have revealed that neurexins also play non-synaptic roles through their regulation of axonal arborization and branching during neural development^50,51^. This conservation of non-synaptic functions for neurexin genes across evolution suggests that neurexin’s involvement in these processes is indeed important for proper neuronal development and function.

In addition to being able to execute dual roles in synaptic function and axonal pathfinding, neurexin genes (and their ligands) have evolved in a manner that makes them well suited for regulating synaptic connectivity. Although there are only three neurexin family members, owing to the presence of multiple promoters and alternative splicing these three neurexin genes can give rise to upwards of 1,000 different neurexin isoforms. Importantly, these different neurexin isoforms have different affinities for neurexin *trans*-synaptic ligands. For instance, LRRTMs and neuroligins exhibit a preference for binding to neurexin variants lacking alternative splicing site 4 (AS4) rather than those containing AS4 ^44,52^. Conversely, cerebellins show a higher affinity for AS4+ neurexin variants compared to AS4-variants^53^, whereas neurexophilins bind specifically to AS2-neurexin variants^54^. The binding affinity of neurexin ligands to the neurexins can similarly be influenced by alternative splicing of the ligands themselves. This has been best demonstrated in the case of neuroligin where the presence of splice site B in neuroligin reduces its binding affinity for alpha and beta neurexins^47^. Together, the diversity of neurexin isoforms and the fact that these different isoforms possess different properties, makes them ideal candidates for regulating synaptic connectivity in the mouse olfactory system, which contains greater than 1,000 different sensory neuron subtypes, each of which must project their axons to a stereotyped location. Consistent with this, our single cell transcriptional analysis has revealed that each of these OSN subtypes expresses a distinct pattern of neurexin expression. Although further work beyond the scope of this current study will be required to fully characterize the expression of all neurexin isoforms in each OSN subtype, the potential richness of diversity of expression of neurexin isoforms is reminiscent of other large gene families such as the DSCAMs and protocadherins, which have been shown to play similarly critical roles in synaptic connectivity in a variety of contexts.

Although our present study has focused on the olfactory system, it is intriguing to speculate that neurexins may play a broad role in regulating synaptic connectivity throughout the brain. Prior work has revealed that neurexin isoforms and their ligands display brain and cell type specific expression profiles suggesting the possibility that these interactions regulate anatomical wiring across the brain during development^55^. Notably, mutations and deletions in neurexin have been associated with several neurodevelopmental disorders, such as ASD and schizophrenia^56–58^. Although at least part of the contribution of neurexin to neurodevelopmental disorders may be explained by their role in synaptic functioning, given that disruption of neuronal connectivity underscores a number of neurodevelopmental disorders^59–63^, further characterizing the role of neurexins and their ligands in regulating anatomical wiring may enhance our understanding of the etiology of these disease states.

## Material & Methods

### Mice

Neurexin1 alpha, 2 alpha, and 3 alpha single knockout mice were derived from B6;129-Nrxn3^tm1Sud^ Nrxn1^tm1Sud^ Nrxn2^tm1Sud^/J mice obtained from Jackson Laboratory (# 006377). Triple neurexins beta conditional knockout mice (#008416), neurexin1 conditional truncation mice (#021777), and neuroligin1 conditional knockout mice (#023646) were obtained from Jackson Laboratory. Triple neurexin conditional knockout mice were generously provided by Takeshi Uemura (Shinshu university). MOR28-IRES-GFP mice were a kind contribution from Gilad Barnea (Brown University) and MOR174-9-IRES-GFP mice were gifted by Bob Datta (Harvard University). M72-IRES-tauCherry mice were obtained from Jackson Laboratory (#029637). E2A-Cre mice was obtained from Jackson Laboratory (#003724). Goofy-Cre mice were gifted by Yoshihiro Yoshihara (RIKEN Center for Brain Science), and Pcdh21-Cre mice were generously provided by Mineto Yokoi (Kyoto University).

All animal care and protocols were followed in accordance with federal guidelines, maintaining a 12-hours light/12-hours dark cycle, 20–23 °C temperature, and 30–70% humidity. These procedures were ethically approved by the University of Massachusetts Medical School Institutional Animal Care and Use Committee (Protocol A-2645-18).

### Plasmids

To obtain purified His-tagged recombinant soluble proteins, the Ntn1-His-Fc plasmid (backbone; pCMVi-SV40ori) was employed, acquired from Woj Wojtowicz (Addgene, #72104), which encodes the signal peptide and extracellular domains of netrin1 fused to the Fc region of human IgG1 along with a C-terminal 6X histidine tag.

In this plasmid, the Ntn1 sequence was replaced with the signal peptide and extracellular domain sequences of neurexin ligand genes, including Nxph1, Nlgn1, Lphn1, Clstn3, and LRRTM2 (Supplementary Figure 4B). Signal peptide and extracellular domain sequences of Nxph1 (NM_00875.5), Lphn1 (NM_181039.2), and Clstn3 (NM_153508.4) were synthesized by GENEWIZ, while sequences of Nlgn1 and LRRTM2 were amplified from pCAG-NL1(-) (Peter Scheiffele, Addgene, #15260) and FSW-HRP-V5-LRRTM2 (Alice Ting, Addgene, #82537) plasmids, respectively. The Ntn1 gene sequence was removed using restriction enzymes, and the synthesized or amplified sequences of neurexin ligand genes were inserted using Gibson Assembly technology^64^. His-Fc, a negative control, was generated by replacing the Ntn1 sequence with only the signal peptide sequence of Nlgn1 in a similar manner. For generating the Nlgn1B-His-Fc plasmid, the splicing site B sequence was incorporated into Nlgn1-His-Fc plasmid using Gibson Assembly.

To generate the HA-tagged neurexin1 alpha, HA-fused Nrxn1α gene was amplified from pCAG-HA-rat Nrxn1 alpha, gifted by Peter Scheiffele (Addgene, #59409), and inserted into pcDNA3.1(+) using Gibson Assembly. The AAV-CMV-GFP plasmid used for neurite outgrowth assay was a gift from Connie Cepko (Addgene, #67634).

### Antibodies

Primary antibodies and their diluted concentrations used were as follows: rabbit anti-mCherry (1:2,000, Abcam, #ab167453), guinea pig anti-VGLUT2 (1:1,000, SYSY, #135 404), mouse anti-His (1:500 for immunostaining and 1:1000 for Western blotting assay, Santa Cruz Biotechnology, #sc-53073), rabbit anti-HA (1:1,000, Cell Signaling Technology, #3724), rabbit anti-Tau (phosphor S396) (1:1,000, Abcam, #ab32057) antibodies.

Secondary antibodies and their diluted concentrations used were as follows: Alpaca anti-rabbit-Alexa488 (1:333, Jackson Immunoresearch, #611-545-215), alpaca anti-rabbit-rhodamine red X (RRX) (1:333, Jackson Immunoresearch, #611-295-215), goat anti-rabbit-Alexa647 (1:333, Invitrogen, #A-21245), goat anti-guinea pig Alexa488 (1:333, Invitrogen, #A11073), goat anti-guinea pig Alexa568 (1:333, Invitrogen, #A11075), goat anti-guinea pig Alexa647 (1:333, Invitrogen, #A21450), HRP-linked horse anti-mouse IgG (1:10,000, Cell Signaling Technology, #7076) antibodies.

### Single-cell RNA sequencing (scRNA-Seq) of mouse olfactory epithelial cells

The previously published 3’ scRNA-Seq data of olfactory epithelial cells was utilized (GSE169021)^65^. Additionally, 5’ scRNA-Seq data of olfactory epithelial cells was newly generated through the following procedure in the current study. For the generation of olfactory epithelial single-cell suspensions, 4 adult male wild-type mice (2 male and 2 female) were euthanized according to our IACUC protocol. The main olfactory epithelia were dissected from the bone under a dissection stereo microscope and quickly immersed in ice-cold Hank’s Balanced Salt Solution (HBSS, Gibco, #14175103). Each epithelium was then washed once with HBSS and resuspended in Earle’s Balanced Salt Solution (EBSS, Worthington, #LK003188) containing papain (20 U/mL, Worthington, #LK003178) and DNase I (2,000 U/mL, Worthington, #LK003172). After a 40-minute incubation with gentle agitation at 37 °C, the suspension was washed twice with phosphate-buffered saline (PBS) containing 0.02% Bovine Serum Albumin (BSA, Worthington, #LS000291) twice and passed through a 30 μm cell strainer (Miltenyi Biotec, #130-098-458).

Single-cell sequencing libraries were prepared using the Chromium Next GEM Single Cell 5’ Reagent Kit V2 (10X Genomics) for sequencing the 5’ end of transcripts, following the manufacturer’s protocols. The libraries were sequenced with 150 cycles of paired-end reads using Illumina Hiseq4000 and Novaseq6000 instruments (Novogene). Sequencing reads were processed through the DolphinNext Single Cell-10X Genomics pipeline (https://dolphinnext.umassmed.edu/index.php?np=1&id=420), employing default settings, with STAR v2.6.1 used for alignment, and the transcriptome build based on gencode M25 with the modification, including the neurexin isoform annotation^66,67^. The OR_deconvolution script (https://github.com/elisadonnard/ORdeconvolution) was applied to correct mis-mapped OR reads^65^.

### scRNA-Seq data analysis

Data analysis was performed using Seurat (v4.9.9 or higher) in R^68^. Cells with fewer than 1,500 unique molecular identifiers (UMIs), more than 50,000 UMIs, fewer than 500 detected genes, more than 1% hemoglobin transcripts, or more than 10% mitochondrial transcripts were excluded. The Scrublet package (0.2.2) in Python (3.7.6 or higher) was utilized to calculate doublet scores, and cells with high doublet scores and/or expressing multiple markers of different cell types were removed. The filtered count matrix was then normalized with log transformation, and the top 2,000 variable genes were chosen for principal component analysis (PCA). Batch effects in PCA space were corrected using Harmony^69^. The number of principal components used for generating UMAP plots was determined using the “Elbowplot” function. Wilcoxon rank-sum test-based differential gene expression analysis (“FindAllMarkers” function) was conducted to identify cell types among the olfactory epithelial cells.

For the confident isolation of the population of OSNs expressing a single OR, the mature OSN cluster was isolated. Cells with fewer than 2,600 UMIs, more than 15,000 UMIs, or expressing other non-canonical olfactory receptors (Taar, Ms4a, V1r, V2r, Fpr, Tas1r, Tas2r) were excluded. The identity of the OSNs was determined by OR transcripts, and only transcriptomes of OSNs with unequivocal OR identities were included in the gene expression-variance and z-score scaled heatmap analyses.

Log-scaled normalized mean expression and standardized variance scores were computed for each gene within the identified OSN transcriptomes. The clusterProfiler (3.18.1) and the org.Mm.eg.db (3.12.0) annotation databases were used for Gene Ontology (GO) term analysis (http://geneontology.org/). The gene lists for synaptic membrane proteins were compiled and consolidated from Syngo, encompassing: 1) integral component of synaptic membrane (GO:0099699), 2) integral component of presynaptic membrane (GO:0099056), 3) integral component of presynaptic active zone membrane (GO:0099059), and 4) integral component of postsynaptic membrane (GO:0099055), 5) integral component of postsynaptic density membrane (GO:0099061), and 6) integral component of postsynaptic specialization membrane (GO:0099060). The gene list for growth cone-related genes (GO:0030426) was obtained from the Mouse Genome Information (MGI) under the Gene Ontology Project (http://geneontology.org/).

### Single-nucleus RNA sequencing (snRNA-Seq) data analysis

The 3’ snRNA-Seq data of olfactory bulb (OB) cells was utilized from the NeMO Archive (http://nemoarchive.org)^70^. The data analysis was conducted using Seurat (v4.9.9 or higher) in R^68^. SoupX (1.6.2 or higher) was applied to remove ambient RNA counts. Cells with fewer than 1,000 UMIs, more than 35,000 UMIs, more than 6,500 detected genes, more than 1% hemoglobin transcripts, or more than 1% mitochondrial transcripts were excluded. The Scrublet package (0.2.2) in Python (3.7.6 or higher) was employed to calculate doublet scores, and cells with high doublet scores and/or expressing multiple markers of different cell types were removed. The filtered count matrix was normalized with log transformation, and the top 2,000 variable genes were selected for PCA. Harmony was used to correct batch effects in PCA space^69^. The number of principal components used for generating UMAP plots was determined using the “Elbowplot” function. Wilcoxon rank-sum test-based differential gene expression analysis (“FindAllMarkers” function) was performed to identify cell types among the OB cells.

### FACS sorting of specific OSN populations from reporter knockin mice

Olfactory epithelial single-cell suspensions were prepared from M72-IRES-GFP, MOR174-9-IRES-GFP, and M72-IRES-tauCherry mice (mixed genders) following the aforementioned protocol. The resulting cell suspension was stained with DAPI and then subjected to FACS to isolate the high-GFP/tauCherry and low-DAPI cell population. Each collected sample contained 5,000-6,000 cells, and three independent samples were generated through separate experimental iterations for each group of mice.

### Embryonic cortical neuron culture

To generate embryonic cortical cell suspensions, mouse embryonic cortical regions at embryonic day 14-16 were dissected under a dissection stereo microscope and placed in ice-cold HBSS. The cortical cells were then collected in EBSS containing papain and DNase I and incubated at 37 °C for 12 minutes with gentle inversion three times every four minutes. The cortical cell suspension was gently triturated using a pipette and subsequently centrifuged at 1000xg at room temperature (RT) for 5 minutes. The resulting cortical cell pellet was resuspended in a mixture of EBSS, albumin-ovomucoid inhibitor (Worthington, #LK003182), and DNase I and carefully layered on top the albumin-ovomucoid inhibitor to create a continuous density gradient. The gradient solution was then centrifuged at 500xg at RT for 5 minutes, and the resulting pellet was resuspended in neuron culture media composed of Neurobasal medium (Gibco, #21103049), serum-free B-27 (Gibco, #17504044), L-glutamine (Gibco, #25030081), and penicillin-streptomycin solution (Corning, #30-002-Cl). Subsequently, the cortical cells were plated in microfluidic axon isolation silicon device (Xona Microfluidics, #snd150) or 12-well plates (Genesee Scientific, #25-106) pre-coated with neuron coating solution I (Sigma-Aldrich, #027-05) overnight at RT. Every three days, 50% of the media was replaced with fresh neuron culture media.

### RNA extraction and cDNA synthesis

For FACS-sorted OSNs expressing M72, MOR174-9, and MOR28, RNA extraction was conducted using the RNeasy Micro Kit (QIAGEN, #74004) in accordance with the manufacturer’s instructions. In the case of embryonic cortical neurons, RNA was extracted from neurons cultured for 3 days *in vitro* (DIV3), with each well containing 600000 cells. The cultured cortical neurons were suspended in TRIzol reagent (Invitrogen, #15596026), vortexed briefly, and chloroform (Sigma-Aldrich, #288306) was added. Centrifugation at 12000xg at 4 °C for 10 minutes facilitated phase separation, with the transparent supernatant collected and mixed with isopropanol (Sigma-Aldrich, #190764). Further centrifugation under the same conditions enriched the RNA pellets, which were then washed three times with 70% EtOH and eluted in Nuclease-free distilled water (Invitrogen, #10977015).

First-strand cDNA was synthesized from the extracted RNA samples using the SuperScript First-Strand Synthesis Kit (Invitrogen, #11904018) following the manufacturer’s instructions.

### Quantitative PCR analysis

Quantitative PCR for gene expression analysis was conducted in 96-well plates (Applied Biosystems, #4346907) using the StepOnePlus 96-well Real-Time PCR System (Applied Biosystems). Reactions with cDNA samples were prepared using a mixture of 2X SYBR Green Master Mix (Applied Biosystems, #4309155) and specific primers designed based on previous studies^71^. The primer sequences used are as follows: Nrxn1α (forward: 5’-TCCTCTTAGACATGGGATCAGG-3’, reverse: 5’-GTGTAGGGAGTGCGTAGTG-3’), Nrxn1β (forward: 5’-TGGCCCTGATCTGGATAGTC-3’, reverse: 5’-AATCTGTCCACCACCTTTGC-3’), Nrxn2α (forward: 5’-GTCAGCAACAACTTCATGGG-3’, reverse: 5’-AGCCACATCCTCACAACG-3’), Nrxn2β (forward: 5’-CCACCACTTCCACAGCAAG-3’, reverse: 5’-CTGGTGTGTGCTGAAGCCTA-3’), Nrxn3α (forward: 5’-GGGAGAACCTGCGAAAGAG-3’, reverse: 5’-ATGAAGCGGAAGGACACATC-3’), Nrxn3β (forward: 5’-CACCACTCTGTGCCTATTTC-3’, reverse: 5’-GGCCAGGTATAGAGGATGA-3’), Gapdh (forward: 5’-CAAAGTTGTCATGGATGACC –3’, reverse: 5’-CCATGGAGAAGGCTGGGG-3’). Neurexin isoform gene expression was normalized to the stably expressed reference gene Gapdh.

### Tissue slice preparation

Postnatal day 10-12 mice were euthanized, and their nasal structures, including the olfactory epithelia and attached OB, were carefully dissected from the skull. The dissected tissue was fixed overnight in 4% paraformaldehyde (PFA, Electron Microscopy Sciences, #15714) in PBS at 4 °C. The following day, the nasal tissues underwent overnight decalcification at 4 °C in 0.45M EDTA in PBS. Subsequently, the nasal tissues were sequentially immersed in solutions of 10%, 20%, and 30% sucrose (Sigma-Aldrich, #S0389) in PBS overnight at 4 °C. Finally, the tissues were embedded in Tissue Freezing Medium (Tissue-Tek, #4583). Cryosections, each with a thickness of 40 microns, were cut onto Superfrost Plus glass slides (VWR #48311-703) and then stored at –80 °C until immunostaining.

### Purification of recombinant proteins

Recombinant protein production was conducted in accordance with the modified previous protocols^44^. Human Embryonic Kidney (HEK) 293 cells were transfected with plasmids encoding His-tagged recombinant proteins as described in the Plasmids section using lipofectamine 2000 (Invitrogen, #11668019) following the manufacturer’s instructions (four 100 mm plates each plasmid). The transfected HEK293 cells were maintained in complete media, consisting of DMEM-high glucose (Sigma-Aldrich, #D5671), supplemented with 10% heat-inactivated fetal bovine serum (R&D Systems, #S12450H), L-glutamine, and penicillin-streptomycin solution, in a 5% CO_2_ humidified tissue culture incubator at 32 °C.

Two days later, media were collected from the cell plates, which were then refilled with fresh complete media, and centrifuged at 1,500xg at 4 °C for 10 minutes to remove cell debris. The collected supernatants were adjusted to 10 mM HEPES and 1 mM EDTA, mixed with protease inhibitor cocktail (Roche, #11836153001), and incubated with protein A agarose beads (Roche, #11134515001) at 4 °C overnight with shaking. On the following day, media were collected again from the remaining HEK293 cells and centrifuged as before. The supernatants were adjusted and mixed with a protease inhibitor cocktail. Protein A agarose beads were collected from the previous reaction by centrifugation and then incubated with the freshly collected supernatants at 4 °C overnight with shaking. Afterward, the samples were washed three times with cold PBS through sequential centrifugation at 900xg at 4 °C for 3 minutes each.

The recombinant protein samples were eluted from the beads using IgG elution buffer (Pierce, #21004) for 5 minutes on ice, centrifuged, and carefully collected from the agarose beads. The protein samples were equilibrated with 0.1M Tris-Cl (pH9.14) and filtered through a 22 μm PVDF syringe filter (Millipore Sigma, #SLGV004SL).

### Microfluidic axon isolation chamber assay

The microfluidic axon isolation device was utilized to investigate axonal responses of cultured neurons to soluble ligands, following both the manufacturer’s instructions (Xona Microfluidics, #SND150) and prior literature references^37^. Cover glasses of size 22×40 mm (VWR, #16004-318) were sterilized with 95% ethanol for an hour, then air-dried overnight at RT. These cover glasses were coated with neuron coating solution I at RT overnight, washed three times with distilled water, and allowed to dry completely. Similarly, the silicon chambers were sterilized with 95% ethanol overnight, followed by washing with distilled water three times and air-drying. The silicon chambers were assembled onto the prepared cover glasses, ensuring no formation of bubbles within the microtunnels, under a ZOE Fluorescent Cell Imager (Bio-Rad). Neuron culture media were introduced into the left and right wells, with an incubation period of 1-2 hours to allow the media to flow through the channels and microtunnels.

Embryonic cortical single-cell suspensions were generated as described above in the Embryonic neuron culture section. Subsequently, the neuron culture media were removed from the devices, and cortical cells were plated into the top left wells (somal compartment) at a density of 130,000 cells in 20 μl per silicon device. After a 1-hour incubation in a 5% CO_2_ incubator at 37 °C, the devices were replenished with fresh neuron culture media in all wells. On the 3rd day post-plating, 50% of the neuron culture media were removed from all wells. Fresh neuron culture media, both with and without the His-tagged recombinant proteins (1 μg for each protein) were then introduced into the top right wells (axonal compartment) and top left wells (somal compartment), respectively. This process was repeated two days later.

Two days following these steps, the silicon chambers were carefully disassembled from the cover glasses. The cortical neurons present on the cover glasses were washed three times with PBS, fixed with 4% PFA in PBS for 20 minutes at RT, and then washed three times with PBS. These neuron samples were subjected to immunostaining.

### Surface binding assay

Cultured HEK293 cells were grown on 12 mm round cover glasses (Azer Scientific, #200121) pre-coated with poly-D-lysine (Gibco, #A3890401) in 24-well plates and then transfected with a plasmid encoding HA-tagged neurexin1 alpha using lipofectamine 2000. The day following transfection, the HEK293 cells were rinsed three times with HEPES bath buffer composed of 150 mM NaCl, 5 mM KCl, 2 mM CaCl_2_, 1 mM MgCl_2_, 10 mM glucose (Sigma-Aldrich, #G8270), and 10 mM HEPES. Subsequently, the cells were exposed to His-tagged recombinant proteins (1 μg for each protein) in HEPES bath buffer for 5 hours at 4 °C while gently shaking. Following the incubation, the cells were thoroughly washed three times for 5 minutes each with the bath buffer, with shaking at RT. Thereafter, the cells were fixed with 4% PFA in PBS for 20 minutes at RT, washed with PBS three times, and then subjected to immunostaining.

### Immunostaining

Cryosections or fixed cultured neurons were treated with a blocking solution composed of 5% normal donkey serum, 0.1% Triton-X100, and Tris-buffered saline (TBS) for 1 hour at RT. Subsequently, the samples were incubated overnight at 4 °C with primary antibodies diluted in the blocking solution (as described above in the Antibody section): Anti-mCherry and anti-VGLUT2 antibodies or anti-VGLUT2 antibodies alone for axon targeting analysis, anti-Tau antibodies for in vitro axon attraction analysis, and anti-His and anti-HA antibodies for surface binding assay. On the next day, the samples were washed three times with TBST (0.1% Triton-X100 in TBS), incubated with secondary antibodies diluted in the blocking solution (1:333) for 1 hour at RT, and then washed three times with TBST. For surface-immunostaining of HEK293 cells, Triton-X 100 was removed from the blocking solution and all washing steps were carried out using TBS instead of TBST.

Afterwards, the samples were counterstained and mounted using Vectashield antifade mounting media with DAPI (Vectashield Laboratories, #H-1900-2). Nail polish was applied to secure the cover glasses, and the slides were subjected to confocal microscopy as detailed in the following procedures.

### Confocal microscopy

Slides were imaged using an LSM 900 Airyscan2 confocal microscope (Zeiss) equipped with a range of objective lenses: 10X/0.45 M27, 20X/0.8 M27, 40X/1.1 water Corr M72, and 63X/1.4 oil DIC. For improved image quality, digital images obtained were refined through the application of a median filter, which effectively eliminated debris significantly smaller than the structures under analysis. Moreover, multi-channel Z-stacks were transformed into two-dimensional representations using Zen blue 3.1 software (Zeiss) to further enhance visualization.

### Analysis of OSN axon targeting

Whole OBs from mice were cryosectioned into 40-micron slices, subjected to immunostaining, and imaged using confocal microscopy with 10X or 20X objectives, as described above. Glomeruli were identified by VGLUT2 signals, and the number of VGLUT2-positive glomeruli, which contained noticeable fraction of axons from specific OR+OSNs as indicated by GFP or mCherry signals, was quantified on both the lateral and medial sides of the OBs. Large glomeruli positive for VGLUT2 and GFP/mCherry were considered correctly positioned original glomeruli, while additional glomeruli containing a smaller fraction of axons from specific OR-positive OSNs were categorized as cases of axonal mistargeting.

### Analysis of microfluidic axon isolation chamber assay

Quantitative assessment of *in vitro* axon attraction was conducted following established literature procedures^37^. For each sample treated with a soluble ligand, neuronal cell bodies of neurons in somal compartment and axons in axonal compartment were sequentially imaged using confocal microscopy with 10X objective (from soma to axonal compartment and from top to bottom of the sample). Using Image J software (National Institute of Health), axonal intensities at 60-micron intervals along the proximal-to-distal axis of axon compartment and cell body intensities in soma compartment were quantified in the combined sequential images. Subsequently, axonal fluorescence intensities were normalized to average neuronal soma intensities. This process was repeated for each set of sequential images collected from top to bottom of each sample, and average intensity was calculated for each ligand-treated sample. Thereafter, the area under the curve (AUC) of the fluorescence intensity graph of axons responding to each ligand was computed for statistical analysis between different ligands.

### Measurement of OB size and glomerular count

OBs were meticulously extracted from 3-4 independent P12 mice per each group, with the attached brain tissue. After a thorough PBS wash, all specimens were fixed overnight at 4 °C with 4% PFA in PBS. The following day, brain and OB samples were triple-rinsed with PBS, photographed using a smartphone camera alongside a ruler for scale (Supplementary Figure 2A), and analyzed for bulb size using Image J software.

Following the measurement of OB size, the bulbs were cryosectioned into 40-micron slices, following the procedure described above. From each mouse, 3-4 sections with consistent positioning within the OBs were selected and subjected to immunostaining using anti-VGLUT2 antibodies. Subsequently, all samples were visualized using confocal microscopy with 10X objective. Using Image J software, no significant difference in the average perimeter of OB sections among the mouse groups was confirmed. Thereafter, the number of glomeruli in each section was quantified.

### Neurite outgrowth assay

Embryonic cortical single-cell suspensions were prepared, as described above in the Embryonic neuron culture section, and seeded onto 18 mm round cover glasses (Azer Scientific, #200181) pre-coated with neuron coating solution I. The cells were plated in 12-well plates with a density of 250,000 per well and cultured in neuron culture media.

Two days later, the cultured neurons were transfected with a GFP-encoding plasmid using calcium phosphate precipitation. Briefly, the plasmids were mixed with 2.5 M calcium chloride in 2X HEPES-buffered saline (50 mM HEPES, 10 mM KCl, 12 mM D-glucose, 280 mM NaCl, and 1.5 mM Na_2_PO_4_, pH 7.06) and the resulting mixtures were added dropwise to each well after a 20-minute incubation. After one hour, the neurons were washed three times with HBSS three times and then incubated at 37 °C and 5% CO_2_. On the same day, 50% of the neuron culture media was replaced with fresh media containing His-tagged recombinant proteins (1 μg for each protein). This step was repeated two days later. Two days after these treatments, the neurons were washed three times with PBS, fixed with 4% PFA in PBS for 20 minutes at RT, washed three times with PBS, counterstained, and mounted using Vectashield antifade mounting media with DAPI (Vectashield Laboratories, #H-1900-2). Nail polish was applied to secure the cover glasses, and the slides were subjected to confocal microscopy as detailed in the following procedures.

Randomly chosen fluorescent neurons were visualized using confocal microscopy with 20X objective. Neurite length and soma diameter were quantified using Image J software, and the length of the longest neurite was then normalized to the diameter of the soma, following established literature protocols^72^.

### Western blot analysis

The concentration of recombinant proteins was determined through a Bradford Assay (Bio-Rad, #5000006) using a NanoDrop One Spectrophotometer (ThermoFisher Scientific). Equal amounts of protein (0.25 µg) from each sample were denatured by boiling in 6X Laemmli buffer (Boston bioproducts, #BP-111R) at 100 °C for 10 minutes and then loaded onto precast protein gels (Bio-Rad, #4561084). The gels were transferred onto PVDF membranes using iBlot PVDF gel transfer (ThermoFisher Scientific). The membranes were then immersed in a blocking solution (5% nonfat milk in TBST) for an hour, followed by incubation with ant-His antibody diluted in the blocking solution overnight at 4 °C with gentle agitation. The following day, the membranes were washed three times with TBST, incubated with HRP-linked horse anti-mouse IgG antibody diluted in the blocking solution for an hour at RT, and then washed three times with TBST. ECL substrates (Bio-Rad, #1705060) were applied to the membrane samples for 5 minutes following the manufacturer’s instructions, and the bands were visualized using the Chemidoc imaging instrument (Bio-Rad).

### Multiplexed error-robust fluorescence *in situ* hybridization (MERFISH) of mouse OBs

MERFISH was performed following the manufacturer’s instructions (Vizgen) and previous literature^36^. Briefly, mice were euthanized, and brains were immediately placed in cold OCT solution. Embedded brains were then frozen at –20 °C and transferred to – 80 °C for storage. Tissue sections of 10-microns were obtained and placed on a functionalized coverslips covered with fluorescent beads. Once adhered to the coverslips, tissues were fixed with 4% PFA in PBS at RT for 15 minutes, followed by three washes with 1X PBS. After aspiration, 70% EtOH was added to permeabilize the tissues for 24 hours. Following a wash with formamide wash buffer (30% formamide in 2X saline sodium citrate (SSC)), the samples were hybridized with a custom MERFISH probe library (Vizgen) for 48 hours. Subsequently, the samples were washed at 47 °C with formamide wash buffer twice, and the tissues were embedded in a polyacrylamide gel, followed by incubation with tissue clearing solution (2XSSC, 2% SDS, 0.5% Triton X-100, and 1:100 protease K) overnight at 37 °C. The tissues were then washed and further hybridized for 15 minutes with the first hybridization buffer, which contained the readout probes associated with the first round of imaging. After washing, the coverslips were assembled into the imaging chamber and placed into the microscope for imaging.

MERFISH imaging was conducted on an automated Vizgen Alpha instrument using imaging buffers, hybridization buffers, and parameter files provided by Vizgen. Initially, a low-resolution mosaic image was acquired (DAPI channel) with a low-magnification objective (X10). Subsequently, the microscope was switched to a high-magnification objective (X60), and seven 1.5 μm z-stack images of each field of view position were generated in 750 nm, 650 nm and 560 nm channels. A single image of the fiducial beads on the surface of the coverslip was acquired and used as a spatial reference. After each round of imaging, the readout probes were extinguished, and the samples were hybridized with another set of readout probes. This process was repeated until combinatorial FISH was completed. Raw data were decoded using the MERLIN pipeline (v.0.1.6, provided by Vizgen) with the codebook for the library used.

### MERFISH data analysis

The Data analysis was conducted using Seurat (v4.9.9 or higher) in R^68^. The count matrix underwent normalization through log transformation, and the top 2,000 variable genes were selected for PCA. Batch effects in PCA space were corrected using Harmony ^69^. The determination of the number of principal components for generating UMAP plots was accomplished using the “Elbowplot” function. The OB layer, olfactory nerve layer, glomeruli layer, mitral cell layer, granule cell layer, and rostral migratory stream were defined based on the expression of Omp, Grem1, Cdhr1, Myo16, and Sox11, respectively.

### Statistics and reproducibility

For the quantitative PCR-based gene expression analysis, three independent samples were used for each group, and at least two separate experiments were conducted. Statistical significance in differences between sample groups was calculated using a post-hoc Tukey’s test or Dunnett’s test following one-way ANOVA. In the case of *in vivo* OSN axon targeting analysis and *in vitro* axon attraction analysis, data were obtained from a minimum of three independent experiments per group. Statistical significance in differences between groups was determined using a post-hoc Fisher’s LSD test following one-way ANOVA or an unpaired t test. For *in vitro* neurite outgrowth analysis, data were collected from three independent experiments per group. Statistical significance in differences between groups was evaluated using a post-hoc Dunnett’s test following one-way ANOVA. In the analysis of OB size and glomerular number, data were gathered from at least three mice for each group. Statistical significance in differences between groups was assessed using a post-hoc Dunnett’s test following one-way ANOVA or unpaired t test.

Further details regarding statistics and sample size can be found in the Figure legends. In this article, the following symbols indicate statistical significance: ****, p<0.0001; ***, p<0.001; **, p<0.01; *, p<0.05; n.s, p ≥ 0.05.

### Data and code availability

OSN scRNA-Seq data described in this manuscript are deposited at GEO accession GSE169021 and GSE248852 (token: mpkzikcmvzuxnob). OB snRNA-Seq data described in this manuscript were obtained from the NeMO archieve (http://nemoarchive.org)^70^. The processed MERFISH data can be found at: https://zenodo.org/records/10420620. All the scripts used for this study can be found at: https://github.com/Greerlab/Nxrn_2023_paper.

## Acknowledgment

We thank Judy Lieberman and members of the Greer Lab for helpful comments on the manuscript. We thank the Flow Cytometry Core for aid in running the FACS experiments. We would like to thank Bob Datta and Gilad Barnea for providing mouse lines. P.L.G. is supported by fellowships from the Smith Family Foundation, the Searle Scholars Program, the Rita Allen Foundation, the Whitehall Foundation, and by grants DP2 OD027719-01 and NIH 5 KL2 TR001455-04 from the National Institutes of Health. D.K is supported by the grant R01GM148832 and K.F is supported by the grant R01MH130582.

## Author contributions

S.J.P conceived the project, and P.L.G supervised it. S.J.P and P.L.G designed the research. S.J.P conducted most of the experiments with assistance from N.L and H.C.J, while I.H.W performed sequencing data analyses. T.U and K.F provided guidance on mouse breeding, and E.M and D.K advised on data interpretation.

## Supplementary Figure legends

**Supplementary Figure 1 related to Figure 1**.

Neurexin gene expression enrichment in OSNs among various OE cells. OSN; olfactory sensory neuron, SUS; sustentacular cell, INP; immediate neural precursor, GBC; globose basal cell, HBC; horizontal basal cell, OEC; olfactory ensheathing cell.

**Supplementary Figure 2 related to Figure 2**.

(A) Representative images and quantitative analysis of the size of OBs in wild-type (WT) and neurexin1, 2, and 3 alpha single knockout (Nrxn1-3α KO) mice (left panels), and control (ctrl) and E2A-Cre-dependent triple neurexin beta conditional knockout (cKO) mice (right panels). n = 6 for WT, 6 for Nrxn1α KO, 8 for Nrxn2α KO, 6 for Nrxn3α KO, 6 for Nrxn123β ctrl, 8 for Nrxn123β cKO. Data are presented as mean ± SEM, Dunnett’s test following one-way ANOVA.

(B) Representative images of glomeruli in WT and Nrxn1-3α single KO mice (left panel), with quantitative analysis of the total number of glomeruli in the OB (right panel). Sections were chosen from each mouse from similar anatomic locations in the OB, and the number of glomeruli stained with anti-VGLUT2 antibodies was counted. n = 14 for WT, 10 for Nrxn1α KO, 9 for Nrxn2α KO, 9 for Nrxn3α KO. All data were collected from three independent experiments. Data are presented as mean ± SEM, Dunnett’s test following one-way ANOVA.

(C) Representative images of glomeruli in Nrxn123β ctrl and E2A-Cre-dependent Nrxn123β cKO mice (left panel), with quantitative analysis of the total number of glomeruli found within the OB (right panel). Sections were chosen from each mouse from similar anatomic locations in the OB, and the number of glomeruli stained with anti-VGLUT2 antibodies was counted. n = 9 for Nrxn123β ctrl, 8 for Nrxn123β cKO. All data were collected from three independent experiments. Data are presented as mean ± SEM, Unpaired t test.

**Supplementary Figure 3 related to Figure 4**.

Z-score scaled heatmap generated by MERFISH displaying the mean expression of neurexin postsynaptic ligands in M/T cell subtypes.

**Supplementary Figure 4 related to Figure 5**.

(A) Quantitative PCR-based normalized expression of neurexin1, 2, and 3 alpha and beta isoforms in DIV3 embryonic cortical neurons. n = 3 for each group. Data are presented as mean ± SEM, ***p < 0.001, ****p < 0.0001, Dunnett’s test following one-way ANOVA.

(B) Schematic diagram of a soluble recombinant protein consisting of signal peptide (SP), extracellular domain of target gene of interest, immunoglobulin Fc region, and 6X His tag (top panel). Analysis of the purified recombinant proteins using Western blotting analysis with anti-His antibody (bottom panel). Red asterisks indicate bands of the expected molecular weights. NLGN1; neuroligin1, NXPH1; neurexophilin1, LPHN1; latrophilin1, CLSTN3; calsyntenin3, LRRTM2; leucine rich-repeat transmembrane protein2.

(C) Surface binding assay performed on HEK293 cells expressing HA-tagged neurexin1 alpha (NRXN1α) using His-tagged recombinant proteins.

(D) Representative images of cultured cortical neurons transfected with GFP exposed to soluble recombinant proteins.

(E) Quantification of the outgrowth of processes in cultured cortical neurons exposed to soluble recombinant proteins. The distance of longest neurite for each cell was normalized to somal diameter. n = 98 for His-Fc, 80 for LRRTM2-His-Fc, 69 for NLGN1-His-Fc, 80 for LPHN1-His-Fc, 73 for CLSNT3-His-Fc. All data were collected from three independent experiments. Data are presented as mean ± SEM, ****p < 0.0001, Dunnett’s test following one-way ANOVA.

