## Supplementary Information for "Combinatorial expression of neurexin genes regulates glomerular targeting by olfactory sensory neurons"

Supplementary Figure 1 related to Figure 1

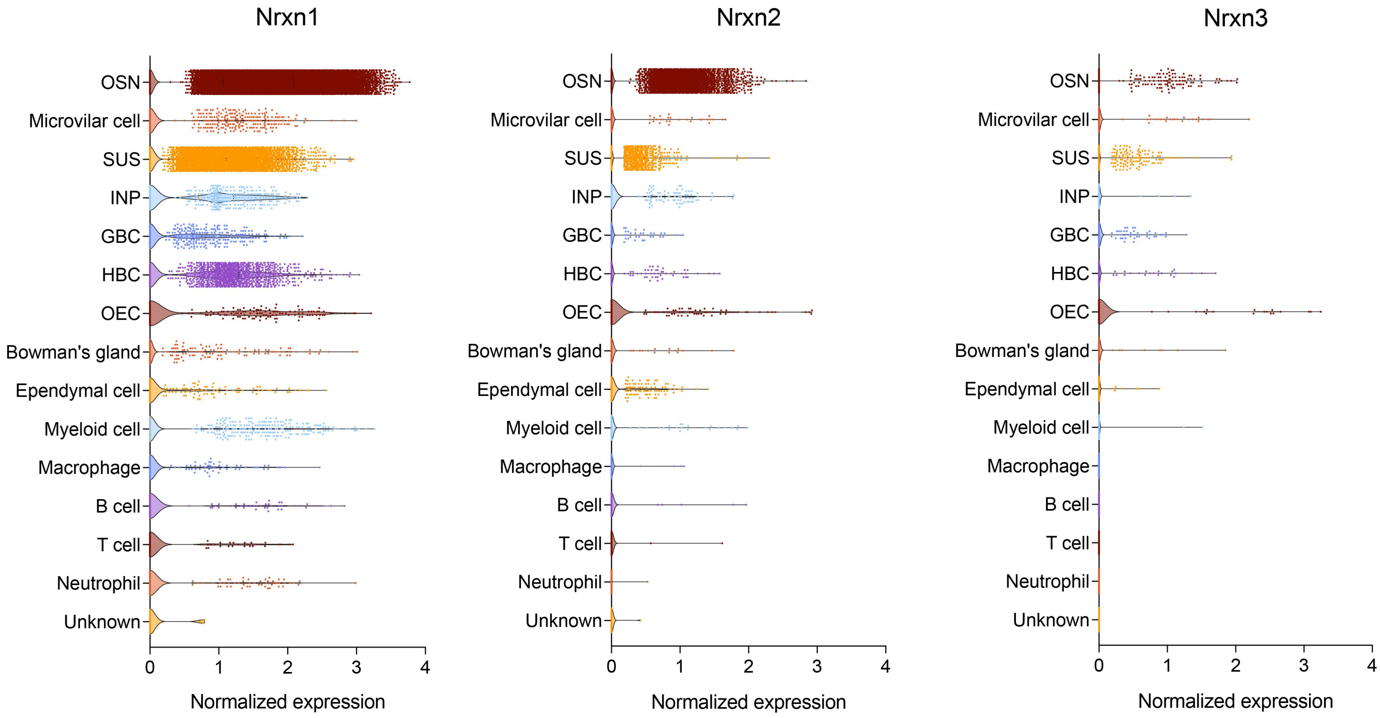

Supplementary Figure 2 related to Figure 2

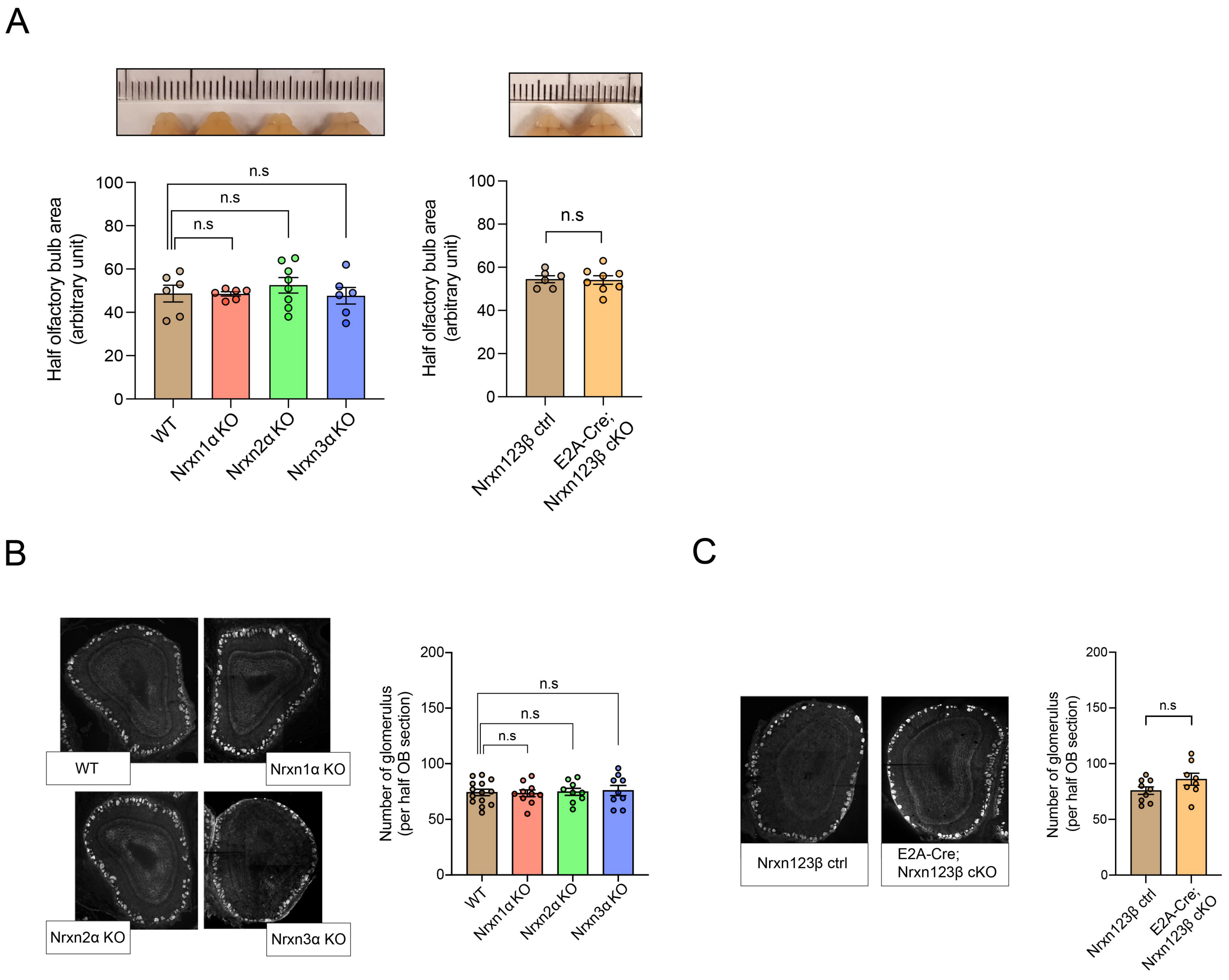

Supplementary Figure 3 related to Figure 4

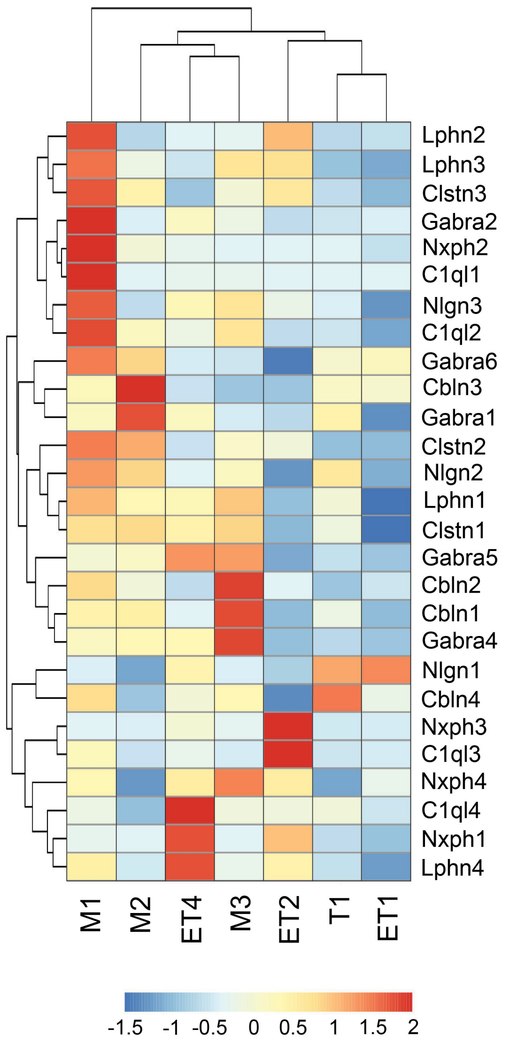

Supplementary Figure 4 related to Figure 5

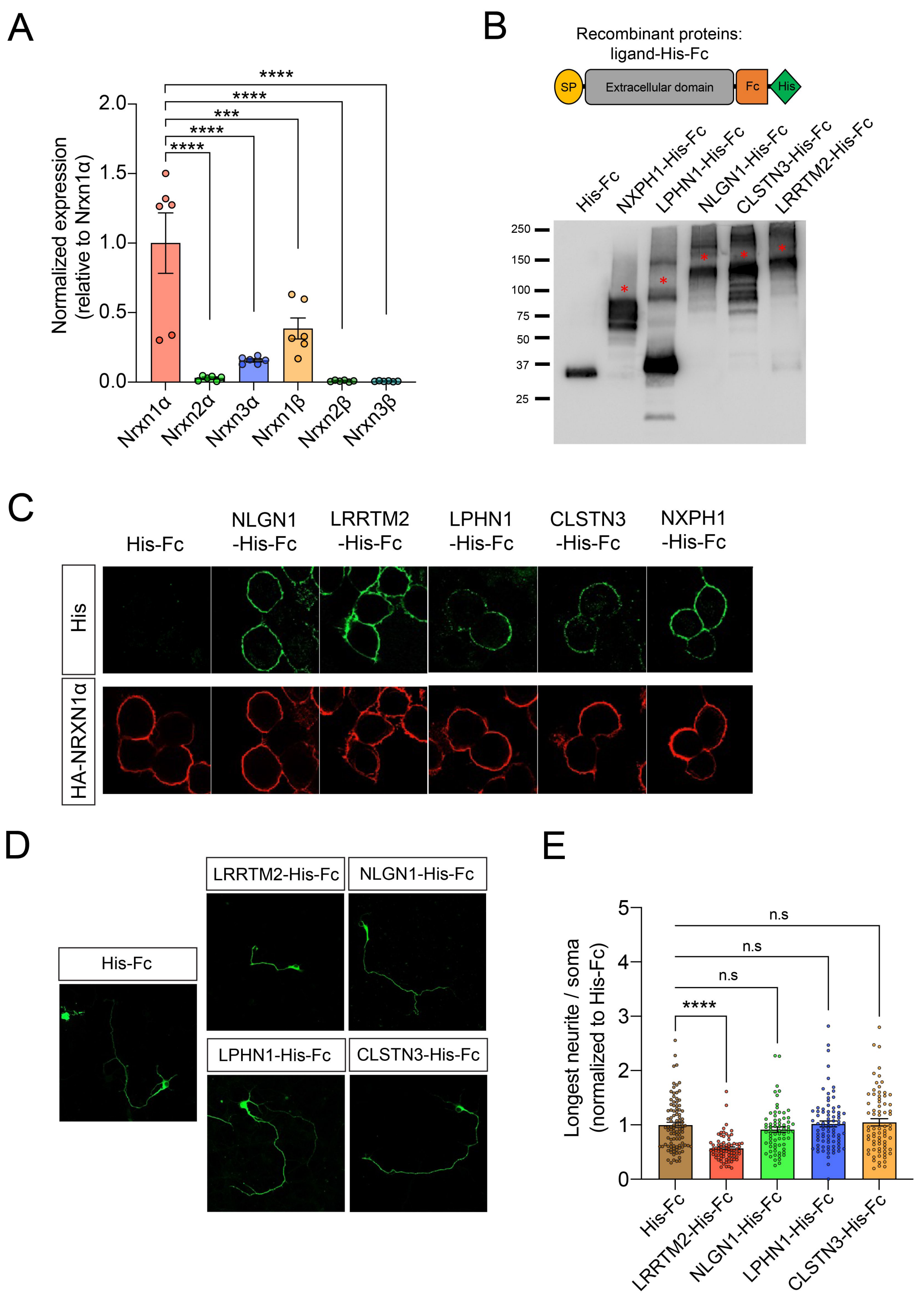
